# Multisite gating in tonic sensory circuits integrates multimodal context to control persistent behavioral states

**DOI:** 10.1101/2022.07.14.500040

**Authors:** Saurabh Thapliyal, Isabel Beets, Dominique A. Glauser

## Abstract

Maintaining or shifting between behavioral states according to context is essential for animals to implement fitness-promoting strategies. How integration of internal state, past experience and sensory inputs orchestrate persistent multidimensional behavior changes remains poorly understood. Here, we show that *C. elegans* integrates food availability and environment temperature over different timescales to engage in persistent dwelling, scanning, global or glocal search strategies matching thermoregulatory and feeding needs. Transition between states, in each case, requires lifting multiple regulatory gates including AFD or FLP tonic sensory neurons activity, neuropeptide expression and downstream circuit responsiveness. State-specific FLP-6 or FLP-5 neuropeptide signaling acts on a distributed set of inhibitory receptors to promote scanning or glocal search, respectively, bypassing dopamine and glutamate-dependent behavioral state control. Multisite gating-dependent behavioral switch by GPCRs in tonic sensory circuits might represent a conserved regulatory logic for persistent behavioral state transitions enabling a flexible prioritization on the valance of multiple inputs.

## INTRODUCTION

Animals continuously integrate information from external environment with their past experience and internal states to generate adaptive behavioral states. This context-dependent modification of behavioral states in turn impacts sensory processing, sensory integration and animal’s response to the environment (Berridge and Waterhouse, 2003; Bramham and Srebro, 1989; Dave et al., 1998). Animals show transient or persistent modifications in the behavioral state based on shift in integrated valance of the context to ensure survival and maximizes their fitness (Gibson et al., 2015; Jo et al., 2020; Sorrells et al., 2022). E.g. animals will use multiple sources of information in order to select between different foraging strategies balancing the risks and benefits of local resource exploitation versus long-range exploration. Failure to execute behavioral state transitions will cause reduced behavioral flexibility, which can impact ecological performance, alter physiology and which is a hallmark of many in human pathologies such as autism spectrum disorders and mood disorders (Devineni and Scaplen, 2021; Lea et al., 2020; Shaw et al., 2002; Uddin, 2021). Recently, several studies in mammals, vertebrates and fruit fly have highlighted function and mechanisms of transition to distinct transient or persistent behavioral states (Andalman et al., 2019; Fu et al., 2022; Jung et al., 2020). However, several questions still remain largely unanswered. (i) How past and current experience of a single cue integrates with context to trigger behavioral state transitions that may persist for hours? (ii) How continuous integration of state-specific signals coordinate multi-dimensional behavioral responses to generate sophisticated and coherent navigation strategy? Due to the relatively complex architecture of their nervous systems, the integrative studies needed to understand the underlying circuit, cellular and molecular mechanisms are challenging in these models.

*C. elegans* has become a popular animal model to understand behavioral state transitions with tools and techniques available to gain multi-layered mechanistic insights (see Flavell et al., 2020 for a review). Previous studies have addressed multiple behavioral states in *C. elegans* based on differential locomotion, egg laying, mate search in response to diverse sensory and physiological cues. In the presence of food, worms show roaming and dwelling states, whereas without food, execute local or global search (Flavell et al., 2013; Fujiwara et al., 2002; Gray et al., 2005; Hills et al., 2004; Shtonda and Avery, 2006). Each of these behavioral states is characterized by a specific locomotory pattern in order to promote a specific exploitation/exploration strategy. E.g., global search combines elevated speed and infrequent turns to increase animal dispersal. Egg laying impacts reproductive fitness and worms display temporal variations in bouts of egg laying activity, which are impacted by feeding status and other sensory cues (Cermak et al., 2020; Waggoner et al., 1998). Sleep and wakefulness are one of the vastly studied behavioral states. Similar to mammals, worms also display sleep-like behavior as ‘lethargus’ during larval stage transitions and ‘sickness-induced sleep’ in response to external physiological stress (Hill et al., 2014; Raizen et al., 2008). Transitions between these diverse behavioral states in response to various environmental cues ensure cellular homeostasis with physiological and reproductive fitness (Skora et al., 2018).

Behavioral states and transitions are characterized based on multi-dimensional phenotypic differences. These are readily quantified with machine-vision-based approach in *C. elegans*, capturing behavioral nuances which are difficult to detect by manual examination and enabling high throughput behavioral state measurement under different conditions (Javer et al., 2018b; Swierczek et al., 2011; Yemini et al., 2013). Moreover, neuron activity recording *in vivo* and manipulation tools allowing mechanistic dissection at circuit levels can link distinct neuronal activity patterns with different behavioral states (Busch et al., 2012; Ji et al., 2021; Kato et al., 2015; Nichols et al., 2017; Venkatachalam et al., 2016). Genetic analyses in different studies revealed the role of conserved neuromodulatory pathways as dopamine, serotonin, tyramine, octopamine and neuropeptide signaling in mediating behavioral state transitions (Bhardwaj et al., 2018; Bhat et al., 2021; Churgin et al., 2017; Flavell et al., 2013; Oranth et al., 2018).

*C. elegans* can sense and modulate behavioral states in response to wide range of sensory and physiological cues such as touch, odors, light, sound, oxygen, CO2, and temperature from the environment (Bargmann, 2006; Bretscher et al., 2008; Ghosh et al., 2021; Goodman and Sengupta, 2019; Gray et al., 2004; Iliff et al., 2021). For our study, we chose temperature as sensory/physiological cue because past and current thermosensory sensory experience can be precisely controlled by varying cultivation temperature and performing temperature shifts. *C. elegans* senses a wide range of temperature, and execute thermotaxis to stay at a preferred temperature in the ‘innocuous range’ of 13-25°C, while avoiding extreme ‘noxious’ temperatures (Aoki and Mori, 2015; Schild and Glauser, 2013; Xiao and Xu, 2021). Based on context, distinct sensory neurons encode temperature information in phasic and tonic neuronal responses and generate appropriate behavioral outputs (Glauser, 2022). Whereas short-lasting processes (second-minutes timescale) underlying temperature-dependent navigation in spatial thermogradient has been extensively studied, we know surprisingly little on temperature-dependent persistent behavioral states and transitions over longer timescales.

Here, we demonstrate that temperature history and current environmental temperature interact with feeding status to drive persistent behavioral changes, to bring animals in two newly described states (*scanning* and *glocal search*, respectively) with unique foraging and thermoregulatory benefits. Each state is controlled by specific neuropeptide-based signaling from two separate tonic thermosensory pathways. In order to compute contextual inputs over time and modalities, a similar logic is used in each pathway, relying on concerted lifting of multiple molecular gates, which occurs only when a specific combination of environmental/internal signals are achieved. Given the conservation of the molecular players involved, we speculate multimodal context processing via multi-site gating mechanisms along tonic sensory pathway might be a widespread regulatory solution mediating persistent behavioral transitions in other species or sensory modalities and its malfunction could underlie human behavioral flexibility disorders.

## RESULTS

### An analytic framework for dissecting temperature and food-dependent behavioral states and transitions

To investigate how multimodal context is processed to orchestrate behavioral state transition and maintenance, we systematically analyzed the behavior of *C. elegans* while varying thermosensory history (growth temperature at 15 versus 25°C), current thermosensory inputs (with thermal shits from 15 to 25 or from 25 to 15°C) and food availability (in fed versus starved animals). We recorded behavioral state snapshots in isothermal environments (Fig. S1A) and compared multiple timepoints to highlight behavioral transition and persistent states. We then performed detailed quantification of *C. elegans* posture and locomotion, focusing on a set of 47 interpretable behavioral parameters (Fig S1C), and visualized behavioral states in a common multiparametric space from a single Principal component analysis (PCA) gathering all the conditions examined in the present study for wild type (Fig 1 A, D, G, J). Below, we sequentially describe the impact of long-term growth temperature, of current thermosensory inputs and the interaction between temperature and food availability.

**Fig. 1.**
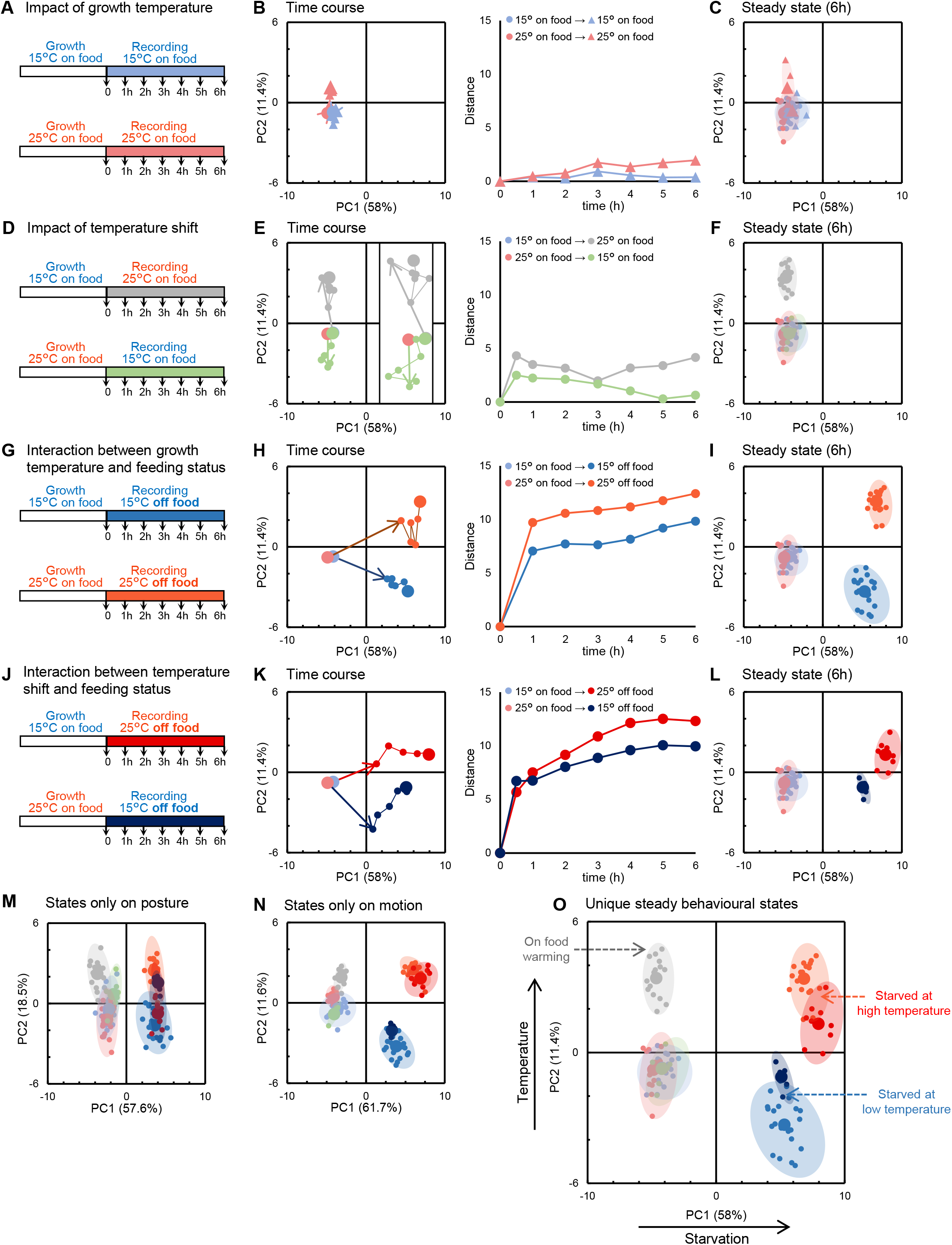
Current and past temperature interact with food availability to set worms in distinct behavioral states. Behavioral states of young adult *C. elegans* exposed to defined thermal and feeding regimen/shifts (as depicted in **A, D, G** and **J**) presented as projections over the two main PCA components from a single analysis of postural and motion parameters over all conditions (**B, C, E, F, H, I, K, L, O**). Behavioral transition unfolding over 6 h presented (i) as its evolution in this PCA space (left panels in **B, E, H, K**, endpoint as larger data mark, first hour changes as arrow) and (ii) as a time courses of the distance between each time point and the starting point (*t=*0) (right panels). Steady states reached after 6 h of the indicated thermal and/or feeding shifts (**C, F, I, L**). Average and individual replicate positions as large and small data marks, resp. 95% CI as colored ellipses. Each replicate as a separate worm population with ≥40 animals. Summary of the four main steady behavioral states adopted by animals in various thermal and feeding contexts (**O**). Similar representations as in **O** for the result of separate PCA focusing on postural (**M**) and motion (**N**) parameters, resp.

### Worms on food are in a dwelling state regardless of their growth temperature

To address the impact of growth temperature, we recorded behavioral snapshots over 6 h of animals cultivated at 15 or 25°C without changing their temperature (Fig 1A). Somewhat surprisingly, we found that animals at different temperatures on food show very similar behavioral states (Lower left quadrant in PCA space, Fig 1B-C). This trend was also confirmed when we separately analyzed the motion and the posture of the animals in two separate PCA analyses (Fig. S1.3 and S1.4). This common behavioral state corresponds to the previously described dwelling state (Flavell et al., 2013; Fujiwara et al., 2002; Shtonda and Avery, 2006; Stern et al., 2017), where animals move slowly and spend most of their time with paused locomotion to feed (Fig 2). Hence, fed animals maintained on food at a constant temperature are in a similar dwelling state regardless of their growth temperature.

**Fig. 2.**
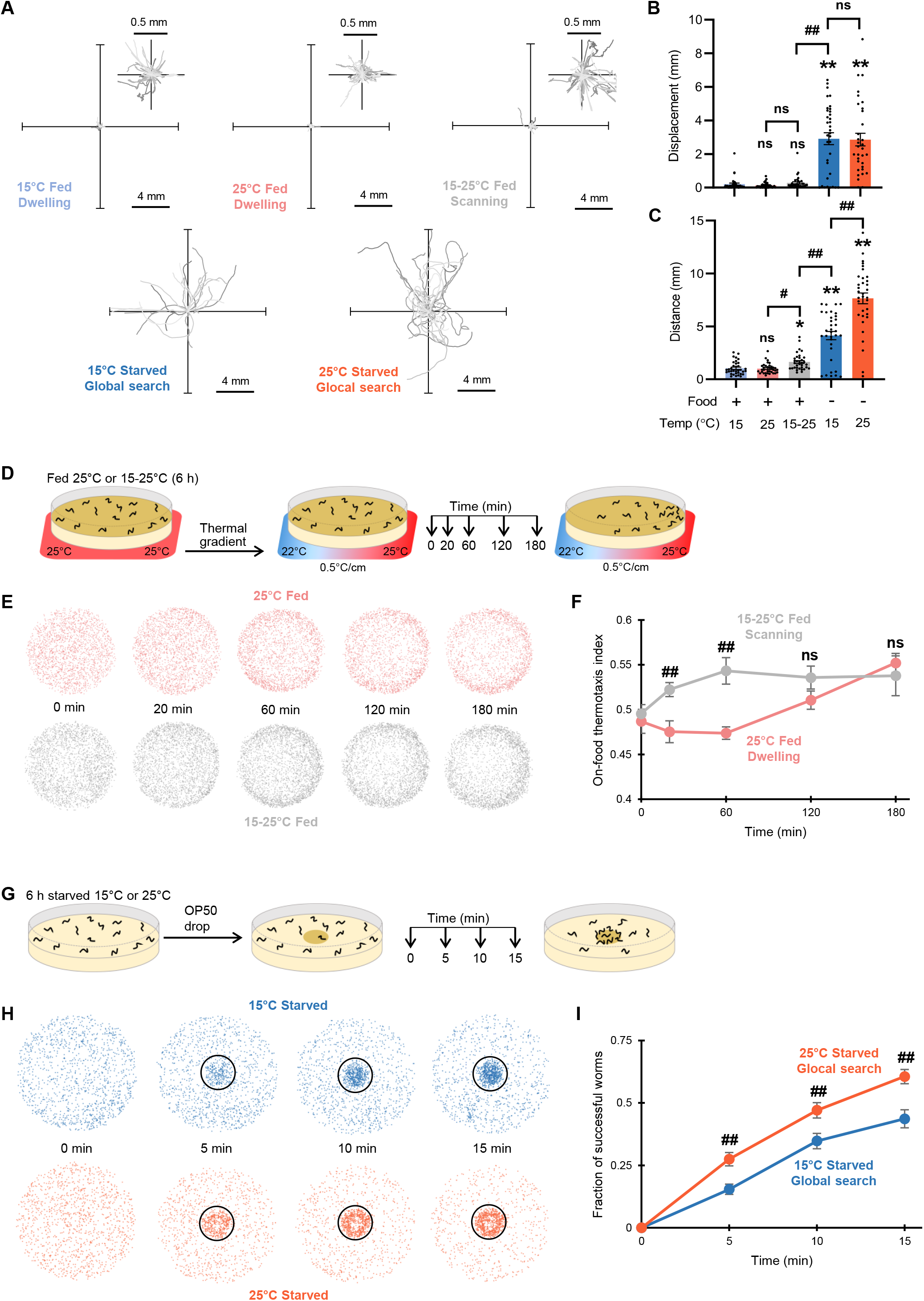
Food and temperature-dependent behavioral states underly different navigation strategies with different food seeking and thermoregulatory performances. One-minute worm trajectories recorded in isothermal environments (35 for each condition) and plotted from a single starting (0,0) coordinate (**A**). Enlarged representation for worms in dwelling and scanning (inset). Average ± s.e.m. and individual data points for animal displacement (**B**, corresponding to how far animals moved from their starting point) and covered distance (**C**, corresponding to the path length). *n*≥30 animals **, *p*<.01 versus 15°C Fed condition, # *p*<.05 and ##, *p*<.01 versus the indicated control by Bonferroni posthoc tests. On-food thermotaxis assay in fed animal revealing a faster thermotactic movement toward recent growth temperature in scanning animals (6h after warming) as compared to dwelling animals held at 25°C (**D, E, F**). Schematic of the assay unfolding (**D**), overlayed worm positions over multiple assays (**E**), on-food thermotaxis index time course (average ± s.e.m., **F**). *n* ≥10 assays each with ≥50 worms. Food drop assay in which starved animals in glocal search mode at 25°C perform better than animals in global search mode at 15°C (**G, H, I**). Schematic of the assay unfolding (**G**), overlayed worm positions over multiple assays (**H**), time course of the fraction of successful worms (average ± s.e.m., **I**). ##, *p*<.01 versus the other condition at the same time point.

### Cooling transiently reinforces dwelling, while warming promotes a persistent scanning sate

Next, to understand how current thermosensory inputs impact behavioral states, we analyzed the behavior of animals grown at 25°C and shifted to 15°C at the onset of the recordings for 6 h (Fig. 1D). In response to cooling, animals underwent a transient transition into a reinforced dwelling state (deeper into the PCA lower left quadrant, Fig. 1E-F), with decreased forward and backward locomotion frequency and concomitantly increased pausing duration and foraging amplitude (Fig. S1.2).

Next, we conducted the reverse experiments and examined the impact of warming, by shifting the animals from 15 to 25°C (Fig. 1D). Animals underwent a long-lasting transition to a new steady state (shift toward the PCA upper left quadrant, Fig1 E-F). Warming modulated both postural (with e.g., reduced tail bending) and motion aspects (Fig. 1M, 1N, S1.3 and S1.4). While keeping low speed when moving, animals increased their foraging speed, reduced pausing time and persistently increased reversals (backward frequency, Fig. S1.2) enabled animals to regularly change their direction. We examined the geometry of 1-min individual trajectories (Fig. 2A) and quantified the displacement of animals (distance from the start and the end of the path, Fig. 2B) and total distance covered (cumulative distance along the path, Fig. 2C). As compared to animals held at 15 or 25°C, animals shifted from 15 to 25 covered more distance, but kept the same low displacement values. Therefore, warming promotes a specific behavioral state, where animals frequently swap between forward and backward motion to more actively scan their local environment while feeding, without dispersing much faster. We will call this new state: scanning.

What could be the advantage of the scanning state? We hypothesized that, in contrast to dwelling animals that barely move, scanning animals may better detect and respond to thermal gradients in order to thermotax while feeding. Indeed, it might be metabolically costly for worm to adapt their physiology to new temperature (Cedergreen et al., 2016) and beneficial to thermotax toward previous growth temperature as a thermoregulatory strategy (Hedgecock and Russell, 1975). We devised an on-food thermotaxis assay in which worms initially held in an isothermal environment at 25°C are exposed to a smoothly appearing thermal gradient (0.5 °C/cm, 22-25°C, Fig 2D). Both animals dwelling at 25 and scanning after warming from 15 to 25°C moved back to 25°C, but the latter drifted much faster (Fig 2E, F).

In summary, shifting worms from 15 to 25°C on food causes them to switch from a dwelling to a scanning state, in which worms adopt a distinct posture and produce short-range movements linked to better thermotactic performances and thermoregulation.

### Starved animals engage in global search at 15°C but glocal search at 25°C

To examine the joint impact of animal feeding status and temperature, we recorded the behavior of starved animals cultivated at 15 and 25°C. In line with the well-documented starvation-evoked shift from dwelling to food-search navigation mode (Gray et al., 2005; Hills et al., 2004; Wakabayashi et al., 2004), starvation at either temperature caused rapid and persistent behavioral changes, affecting both postural and motion parameters (Fig. 1G-I, 3A, A1.3 and S1.4) and causing a right shift in our PCA space (with increasing PC1 value, Fig. 3B). Regardless of temperature, starvation produced a rapid decrease in pausing time and dwelling, with concomitant increase in the time spent in forward locomotion, as well as a progressive increase in reversal duration and postural alterations (Fig. S1.2), consistent with a global search state.

**Fig. 3.**
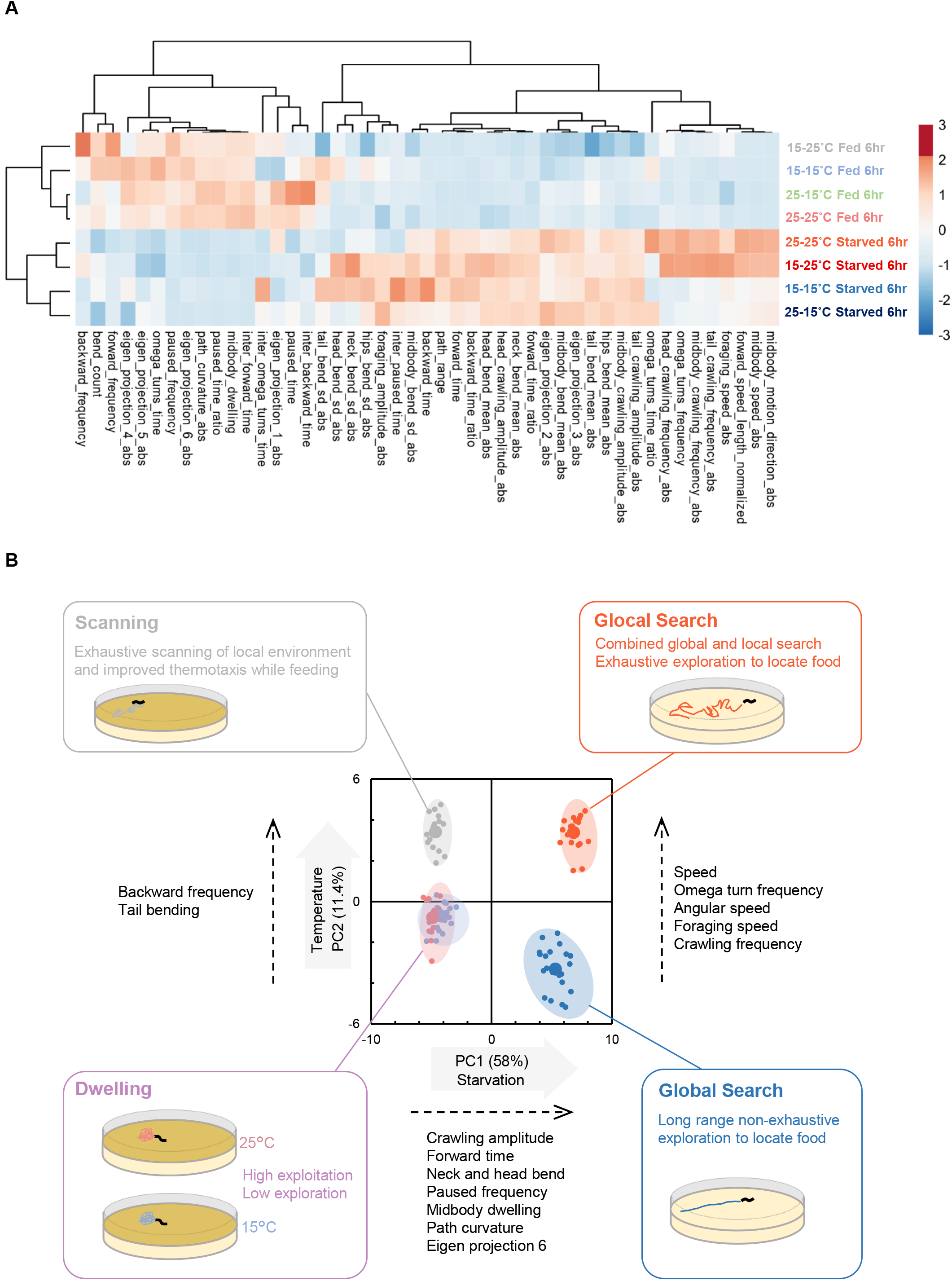
Distinct food and temperature-dependent behavioral codes underlie specific behavioral exploitation/exploration strategies. Behavioral steady-states after 6 h in the indicated food and temperature-dependent condition. Heat-map of behavioral parameters of *z*-scores across the indicated conditions and hierarchical clustering trees based on Euclidian distances (**A**). 25-15 (cooling), 15-25 (warming), and 15-15 or 25-25 (temperature left unchanged). Behavioral state representation as in Fig. 1O, with behavioral state interpretation annotations and illustration of the corresponding exploitation/exploration strategies (**B**).

Very interestingly, however, current temperature had a strong impact to steer worms into distinct behavioral states at 15 or 25°C (lower and upper right quadrants in PCA space, Fig. 1I-L, S1.3). Distinctively after starvation at 25°C, worms increased speed, foraging speed, angular speed, omega turn and crawling frequency and produced longer and more twisty trajectories, contrasting with the shallowly curving radial trajectories at 15°C (Fig. 2A). The worm displacement at 25°C was similar to that at 15°C, however, indicating that the increased locomotory activity at 25°C did not enhanced dispersal (Fig. 2B-C). We obtained similar results with Monte-Carlo simulations emulating worm dispersal with empirically measured speed and turn frequency values (Fig. S2). These observations are consistent with a model in which starved worms at 15°C are in a global search state, dispersing relatively fast to find food without exhaustively sampling the local area along their path, whereas worms at 25°C disperse equally fast, but cover a larger area around their path with concomitant elevation of speed and omega turns. Therefore, starved worms at 25°C are in a state enabling both local and global search, which we will call glocal search.

In addition to increasing the area covered during exploration, we further hypothesized that fast moving animals in glocal search state would more efficiently detect and reach food sources in their surroundings. As an empirical test, we designed an assay in which a food drop was added onto a plate of searching worms and their progression monitored (Fig. 2G). We found that animals in glocal search state at 25°C performed better than animals in global search state at 15°C (Fig. 2H-I).

Collectively, our data indicate that, based on current temperature, starved worms operate multidimensional behavioral changes to select between global or glocal search states, which tunes the efficacy and exhaustiveness of food-seeking exploration.

### Gating mechanisms prevent behavioral transitions except in specific multimodal contexts

As a whole, the analyses reported so far provide a comprehensive picture of how food and temperature-associated contexts can steer worms into four distinct steady behavioral states (separate quadrants in our PCA analysis), each corresponding to a exploitation/exploration strategy with its respective advantages for environment sampling and navigation (Fig. 3B). Whereas in our PCA analysis, PC1 mostly reflects the impact of food availability and PC2 the impact of temperature, these two factors do not produce simple additive effects, but rather interact, which opens new questions. What gating mechanisms prevent scanning entry on food unless a recent warming occurred? How temperature impact is ungated upon starvation for animals to commit into global or glocal search? Below, we dissect the cellular and molecular gating/ungating mechanisms controlling these specific behavioral states.

### Scanning is controlled by AFD, which encodes recent warming as tonic calcium signals

To identify neurons required for warming-evoked transition to scanning on food, we genetically ablated candidate thermosensory neurons (AFD, FLP, AWC and ASI) and quantified backward frequency and tail bending, which are two hallmarks of scanning (Fig 4A and S4A). Changes in both parameters were still observed when ablating AWC, ASI or FLP, but strongly inhibited by the ablation of AFD (Fig 4B and S4B), suggesting a major role for AFD in the persistent behavioral switch to scanning. Blocking synaptic transmission from AFD using Tetanus toxin also abolished the backward frequency and tail bending modulation (Fig 4B and S4B), confirming the importance of neurotransmission from AFD for warming-evoked scanning entry.

**Fig. 4.**
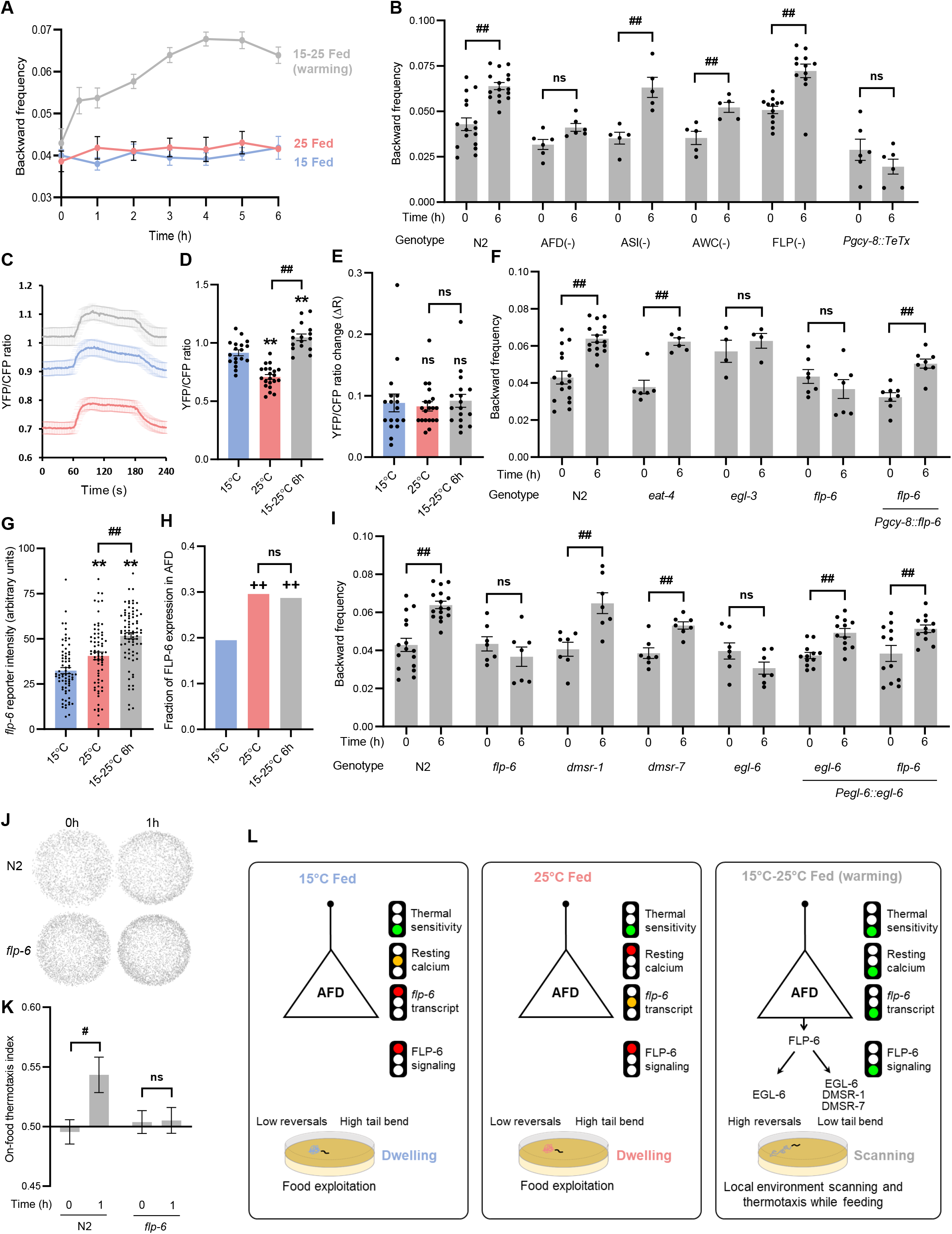
Multisite ungating in AFD thermosensory neurons controls FLP-6-dependent scanning entry. Time course of worm backward frequency increase after warming from 15 to 25°C (**A**). Backward frequency at *t*=0 and *t=*6 h following warming in the indicated genotypes (**B, F, I**). Mean ± s.e.m. of *n*≥4 assays each with ≥50 worms. *Pgcy-8::TeTx*, transgene blocking synaptic transmission in AFD (**B**); *Pgcy-8::flp-6*, transgene for AFD-specific *flp-6* rescue (**F**); *Pegl-6::egl-6*, transgene for *egl-6* over-expression and *flp-6* mutation by-pass analyses in *egl-6* and *flp-6* mutant background, respectively (**I**). Cameleon YFP/CFP ratio in AFD reporting the absolute intracellular calcium levels at rest and following a 2-min 10°C thermal up-step in animals grown at 15°C, grown at 25°C or shifted from 15 to 25°C for 6 h (**C-E**). Mean ± s.e.m. of *n≥*15 traces (**C**), resting calcium levels (**D**) and temperature-evoked relative calcium changes (**E**). *flp-6* transcriptional reporter quantification comparing mean intensity (**G**) and fraction of AFD neurons with detectable signal (**H**). Result of on-food thermotaxis assays after 1 h as in Fig. 2E-F, showing impaired temperature-dependent drift in *flp-6* mutants (**J**-**K**). **, *p*<.01 versus 15°C Fed condition, # *p*<.05 and ##, *p*<.01 versus the indicated control by Bonferroni posthoc tests and ++, *p*<.01 versus 15°C Fed condition by Fisher’s exact test for contingency comparisons. ns, not significant. Schematic of the multisite gating model controlling scanning entry (with gates as traffic lights, **L**). In dwelling animals, at least one gate is closed (red light), whereas they are all lifted (green lights) after warming to promote scanning entry.

AFD neurons can encode thermal history as absolute intracellular calcium levels and simultaneously respond to short-lasting thermosensory inputs with fast-adapting calcium transients (Glauser, 2022; Hawk et al., 2018; Ippolito et al., 2021). To investigate if AFD calcium dynamics could encode information of behavioral state transition, we imaged AFD calcium levels using a ratiometric YC2.3 calcium sensor and compared animals grown at 15°, grown at 25° or exposed to warming (15 to 25°C shift). We measured resting calcium levels (baseline at constant temperature, Fig. 4D) and thermal responsiveness to short-lasting 10 °C thermal up-steps (Fig 4E). Consistent with previous studies, we observed that AFD (i) produced rapid calcium transients in response to thermal up-steps, regardless of growth temperature (Fig. 4C, E) and (ii) showed higher resting calcium levels in animals grown at 15°C compared to 25°C (Fig. 4C, D). A 6 h shift from 15 to 25°C did not alter the magnitude of AFD responses to short-lasting stimuli (Fig 4C, E), but significantly increased the resting calcium level as compared to animals constantly kept at 15 or 25°C (Fig. 5C). These results highlight that absolute calcium level in AFD is tonically modulated over an hour timescale and that this level is particularly high in animals in the scanning state 6 h after warming.

**Fig. 5.**
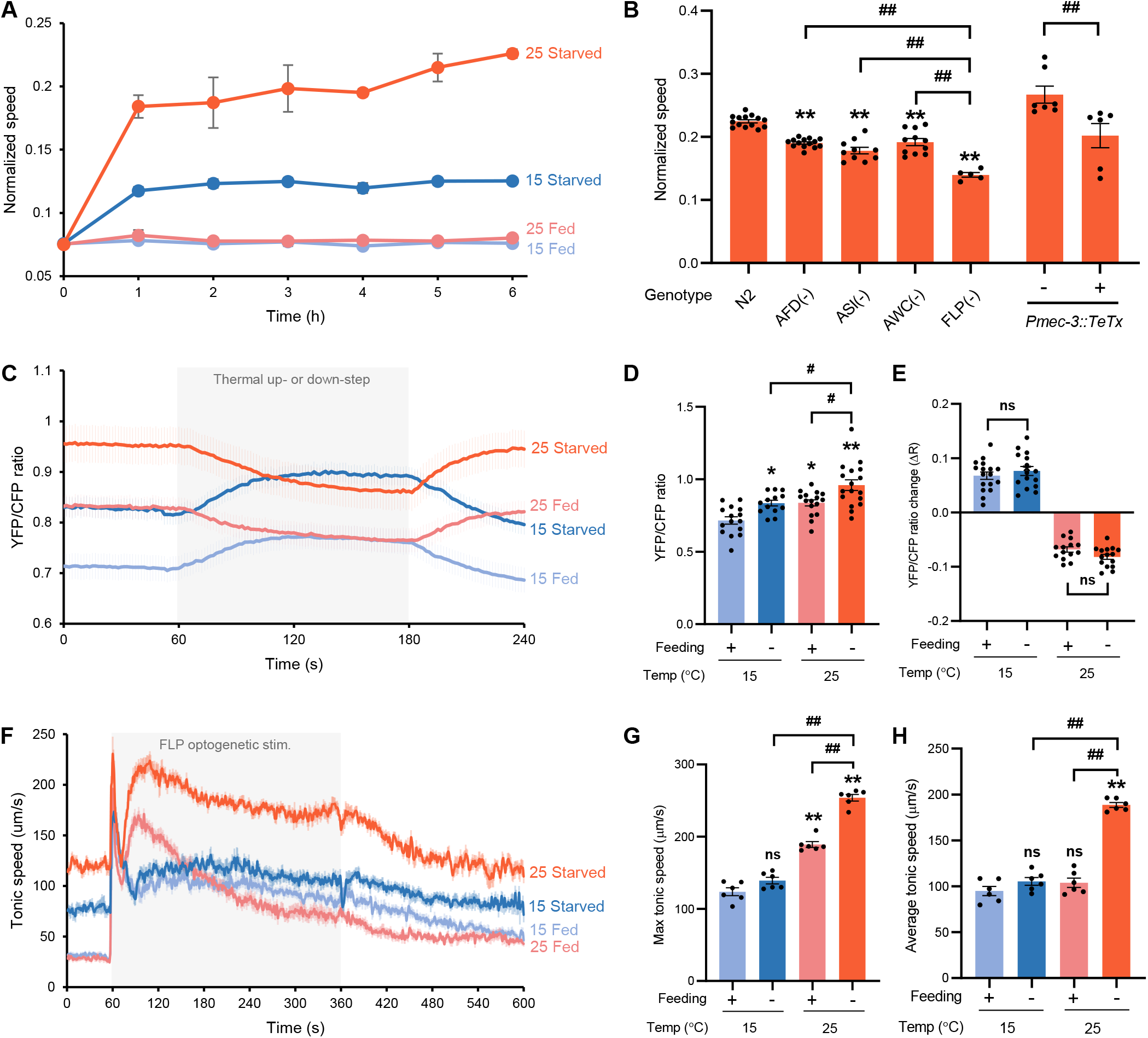
Temperature and feeding states are encoded in FLP neural pathway to control tonic speed elevation during glocal search. Speed increase time course after starvation at 15 or 25°C and in corresponding controls left on food (**A**). Mean ± s.e.m. of *n*=6 assays, each with ≥50 worms. Speed measured after 6h of starvation at 25°C in wild type (N2), in transgenic lines with ablated neurons, or in animals carrying a *Pmec-3::TeTx* transgene blocking neurotransmission in FLP (**B**). FLP absolute intracellular calcium levels at rest and following 2-min 10°C thermal up- or down-steps (**C**-**E**), reported as in Fig. 4C-E and showing FLP encodes temperature and food signals as resting calcium levels without modulating thermal responses to short stimuli. Impact of temperature and food on speed elevations caused by a 5-min tonic optogenetic activation of FLP (**F**-**H**). Mean speed elevation profiles (± s.e.m. of *n=* 6 assays, each scoring *≥*30 worms) with a first spike corresponding to an initial high reversal period (*t*=60-90 s) and a second tonic elevation period (90-360 s) (**F**). Max (**G**) and average speed (**H**) during the tonic speed elevation period. *, *p*<.05 and **, *p*<.01 versus 15°C Fed condition; # *p*<.05 and ##, *p*<.01 versus the indicated condition by Bonferroni posthoc tests. ns, not significant.

### Warming up-regulates *flp-6* neuropeptide expression in AFD to promote scanning and enhance thermotaxis on food

Since the transition to scanning after warming is linked to a tonic AFD activation and requires neurotransmission, we next wanted to identify the relevant communication molecules released by AFD. AFD produces glutamate and several neuropeptides (Ohnishi et al., 2011; Taylor et al., 2021). We thus tested if warming-evoked scanning entry was affected in *eat-4* mutants with impaired glutamate signaling (Lee et al., 1999) and *egl-3* mutants with impaired neuropeptide production (Husson et al., 2006). *eat-4* mutation had no impact, but *egl-3* prevented backward frequency increase and tail bending decrease in response to 6 h warming (Fig. 4F and S4C). These results suggest that neuropeptide not glutamate signaling is required to shift from dwelling to scanning after warming. To uncover the neuropeptide signal, we tested *flp-6* mutants. We chose this candidate because it is the most abundantly expressed neuropeptide gene in AFD (Taylor et al., 2021), its transcription is regulated by temperature and its tonic release mediates temperature-dependent regulation on life span by AFD (Chen et al., 2016). We found that *flp-6* was required to shift both backward frequency and tail bending in response to warming (Fig 4F and S4C). This effect was significantly rescued by a *[Pgcy-8::flp-6]* transgene selectively restoring *flp-*6 expression in AFD (Fig. 4F, S4C). While not ruling out additional mechanisms, these data suggest that FLP-6 signal originating from AFD makes an important contribution to the warming-evoked transition to scanning.

Because FLP-6 neuropeptide is essential to modify key parameters responsible for the scanning state, which increase thermotactic performances, we then predicted that *flp-6* mutants would not perform well in locating and responding to thermogradient on food. According to our prediction, we found that, even after 1 h in a gradient, *flp-6* mutants failed to navigate to warmer side of the plate while N2 navigate within 20 min (Fig. 2F and 4J, K). These data show that the behavioral disruption caused by the lack of FLP-6 alters scanning execution and ensuing thermoregulatory benefits.

Since temperature was previously shown to affect *flp-6* transcription in AFD (Chen et al., 2016), we next used a transcriptional reporter to test if *flp-6* transcription was actually increased in animals in the warming-evoked scanning state. We found that the fraction of animal with detectable *flp-6* reporter signal was significantly higher in animals raised at 25°C or shifted to 25°C as compared to animal raised at 15°C (Fig. 4H). Furthermore, the signal intensity was significantly stronger after the 15-25°C shift (Fig. 4G).

Collectively, our results suggest that warming-evoked transition to scanning on food is controlled by AFD and FLP-6-dependent signaling. A thermal shift from 15 to 25 is linked to a unique AFD state that combines elevated FLP-6 expression and tonic AFD activity concomitant to chronically elevated calcium levels (Fig. 4L). Reduced calcium and low *flp-6* expression represent potential gating mechanism (red traffic lights in the model in Fig. 4L) preventing scanning entry at constant temperature. This multisite gating will only be lifted in response to recent warming, in order for AFD to signal via FLP-6 and promote scanning only when this specific behavioral state is needed to jointly reach feeding and thermoregulation goals.

### Different FLP-6 receptors regulate reversal and tail bending during scanning

*C. elegans* neuropeptides function through GPCRs to generate neuromodulatory effects lasting across timescales. To identify receptors mediating FLP-6-dependent scanning entry *in vivo*, we examined the impact of the loss of *egl-6, dmsr-1* and *dmsr-7* receptor genes, coding for FLP-6 candidate receptors identified in an *in vitro* ligand-receptor screen (Beets et al. in prep.). We found that *egl-6* not *dmsr-1* or *dmsr-7* mutants failed to increase backward frequency, while all three mutants failed to decrease tail bending (Fig. 4I, S4D). We further rescued defects in *egl-6* mutants by expressing EGL-6 under its native promotor. Bypassing the lack of *flp-6* by overexpressing EGL-6 under its own promoter in *flp-6* mutants was sufficient to restore their backward frequency, but not their tail bending phenotype (Fig. 4I and S4D). Taken together, these results suggest that FLP-6 signaling might use a distributed set of receptors to orchestrate the multi-dimensional behavioral changes linked to scanning.

### FLP thermosensory neurons are essential for glocal search

Our systematic characterization of food and temperature-dependent behavioral states (Fig. 1-3) showed that the temperature influence is gated on food to maintain dwelling, but that temperature strongly impacts the choice between global and glocal search. Since speed and omega turn up-regulation are distinctive characteristics of glocal search (Fig. S1.2, 5A and S5A) and suffice to recapitulate the main navigational features of this exploration strategy in simulations (Fig. S2), we focus on these two parameters for further analyses.

To identify neurons required for the transition to glocal search at 25°C, we genetically ablated candidate thermosensory neurons (Fig. 5B and S5B). Ablation of FLP neurons caused severe, while ASI ablation caused moderate impairment of both speed and omega turns. AFD and AWC ablation caused moderate impairment of only speed not omega turn frequency. Blocking synaptic transmission with Tetanus toxin expression in FLP also prevented the elevation of both speed and omega turn frequency. These results highlight a major contribution of FLP neurons in shifting speed and omega turns for transition to glocal search.

### Food and temperature affect tonic calcium activity in FLP and downstream circuit responsiveness

FLPs are non-adapting phasic and tonic thermosensory neurons (Saro et al., 2020). To examine if FLP neurons could jointly encode food and thermal information, we performed *in vivo* calcium imaging of FLP using the ratiometric YC2.3 sensor. Consistent with our previous work (Saro et al., 2020), we found that FLP encodes information of absolute temperature (15°C and 25°C) as resting baseline calcium levels (Fig. 5C, D). Starvation also increased resting calcium in FLP, having an additive effect with temperature. Maximal levels were thus observed in animals in glocal search state after starvation at 25°C. In contrast, we found that responses of FLP to 2-min, 10°C up- or down-steps were neither affect by starvation, nor by temperature (Fig. 5C, E). These results indicate that starvation and temperature independently modulate resting calcium levels (not thermal responsiveness) of FLP neurons, which could partially encode information of distinct behavioral states.

To examine if the responsiveness of components downstream of FLP depolarization are modulated in specific states, we monitored the changes in speed evoked by the optogenetic activation of FLP as a function of temperature and feeding state. Consistent with our previous work (performed in animals starved at 23°C, (Saro et al., 2020)), we found that optogenetic FLP activation first triggered an abrupt speed change associated with a high reversal phase (60-90s), which was followed by a tonic speed induction phase (90-360s) persisting throughout the stimulation (Fig. 6F). Both fed and starved animals at 15°C produced a relatively low tonic speed elevation with a slow decay after reaching max speed (Fig. 6F and G). Fed animals at 25°C produced a more pronounced increase compared to animals at 15°C (Fig. 6F and 6G), but this speed elevation in the tonic phase underwent fast decay (resulting in low average tonic speed measures Fig. 6H). Starved animals at 25°C showed further increase in maximum tonic speed compared to animals on food with minimal decay (Fig. 6F and 6G), resulting in elevated average tonic speed (Fig. 6H).

**Figure 6.**
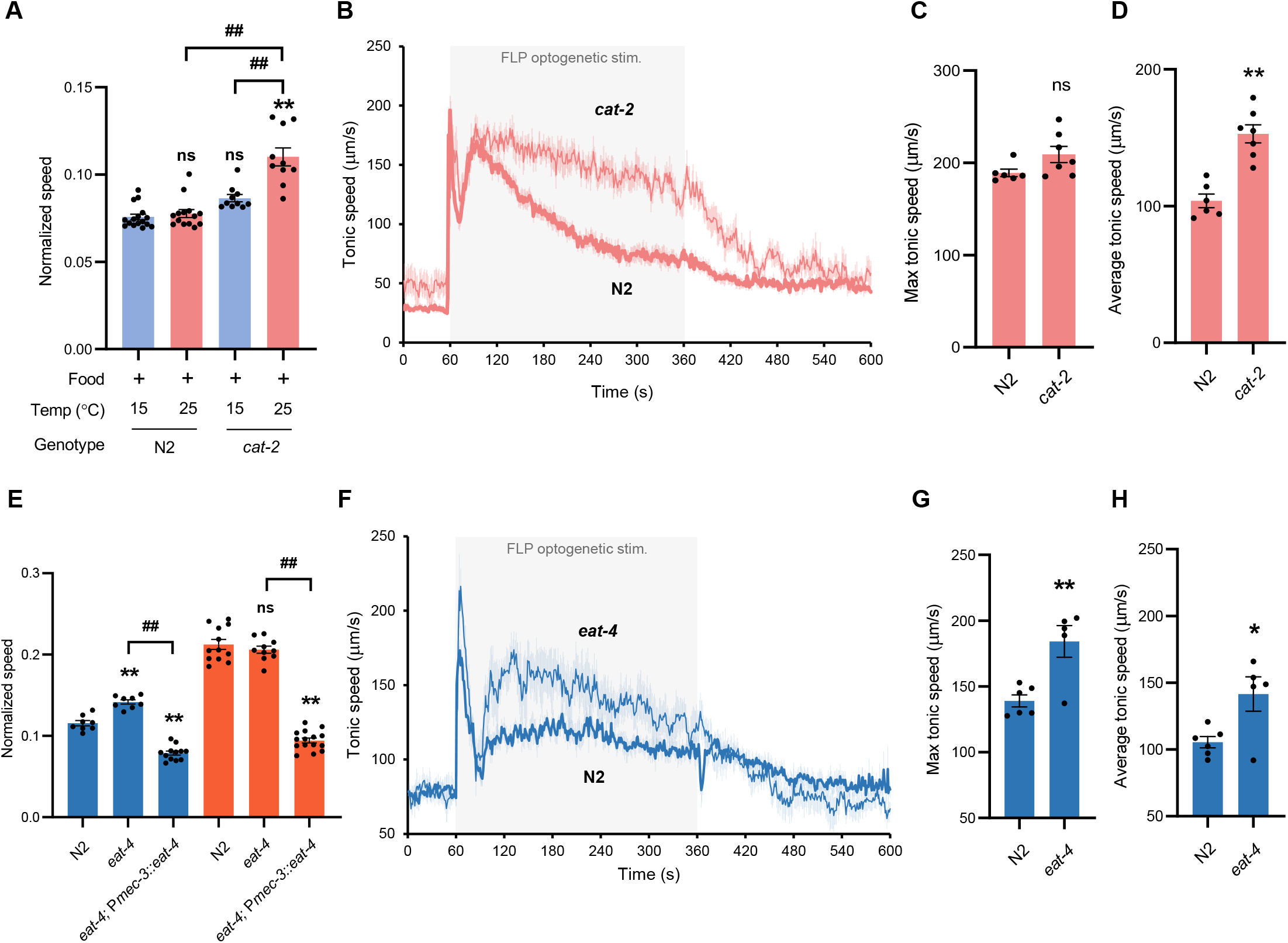
Speed elevation is inhibited by dopamine in fed animals at 25°C and by glutamate signaling in starved animals at 15°C. Impact of a *cat-*2 mutation blocking dopamine biosynthesis (**A**-**D**) on the speed of fed animals at 15 or 25°C (**A**), and on the tonic speed response evoked by a 5-min optogenetic activation of FLP in animal maintained at 25°C (**B**-**D** scored and reported as in Fig. 5). Similar analyses for an *eat-4* mutation affecting glutamatergic signaling in starved animals at 15°C (blue) or 25°C (orange) (**E**-**H**). Data as average ± s.e.m. of *n≥* 5 assays, each scoring *≥*30 worms. *, *p*<.05 and **, *p*<.01 versus 15°C Fed condition or respective N2 control; # *p*<.05 and ##, *p*<.01 versus the indicated condition by Bonferroni posthoc tests. ns, not significant.

Collectively, results from calcium imaging and optogenetic analyses suggest a model in which the temperature and starvation-dependent sustained speed increase underlying glocal search could result from a concomitant increase in FLP tonic activity and enhanced responsiveness of downstream components. In corollary, the lower speed observed in dwelling and global search states seems to result from the gating of the FLP-dependent speed-promoting pathway by food and low temperature.

### Dopamine prevents search behavior on food at high temperature

Dopamine signaling was shown to modulate food-dependent behaviors (Cermak et al., 2020; Hill et al., 2014; Oranth et al., 2018). Next, we tested if dopamine signaling gates temperature-dependent behaviors on food. Unlike wild type, *cat-2* mutants defective for dopamine synthesis had increased speed and omega turn frequency at 25°C compared to 15°C even on food (Fig. 6A and S6A).

Moreover, the fast speed decay observed during FLP optogenetic stimulation at 25°C on food was also revoked in *cat-2* mutants (Fig. 6B-D). These results suggest that active dopamine signaling on food normally inhibits high temperature-evoked behaviors and functions as an important food gate in the context of the transition from dwelling to search behaviors.

### Glutamate signal from FLP hinders speed increase during global search at 15°C

Next, we hypothesized that one or more signals from FLP could modulate speed and omega turns after starvation, keeping in mind that such signals could either act by inhibiting the response at 15°C or stimulating it at 25°C. Since FLP produces glutamate, we first examine starved *eat-4* mutants and found they displayed increased speed at 15°C (Fig. 6E). A *[Pmec-3::eat-4]* transgene restoring *eat-4* expression in FLP neurons could revert the speed to values that were even below those in wild type, potentially due to *eat-4* over-expression (Fig. 6E). The *eat-4* mutation also potentiated the tonic speed elevation during optogenetic stimulation of FLP neurons, with an increase in max and average tonic speed at 15°C (Fig. 6F-H). These results suggest that glutamate signaling in the FLP-dependent neuronal pathway functions as temperature-dependent inhibitory drive for speed after starvation and might be relevant to prevent maximal speed elevation at 15°C, staying at an appropriate level for global search.

### FLP-5 and other neuropeptides from FLP promotes glocal search

Next, we analyzed the contribution of neuropeptide-based signaling. *egl-3* mutation prevented both the increase in speed (Fig. 7A) and in omega turn frequency (Fig. S7A) after starvation at 25°C, indicating requirement of neuropeptide signaling. We performed a biased screen for neuropeptide-encoding genes expressed in FLP neurons and found that the up-regulation of speed requires *flp-5*, while that of omega turns requires *flp-5, flp-13* and *nlp-14* (Fig. 7A and S7A). Both *egl-3* and *flp-5* mutations significantly impaired the tonic speed elevation evoked by FLP optogenetic stimulation (Fig. 7B-D).

**Figure 7.**
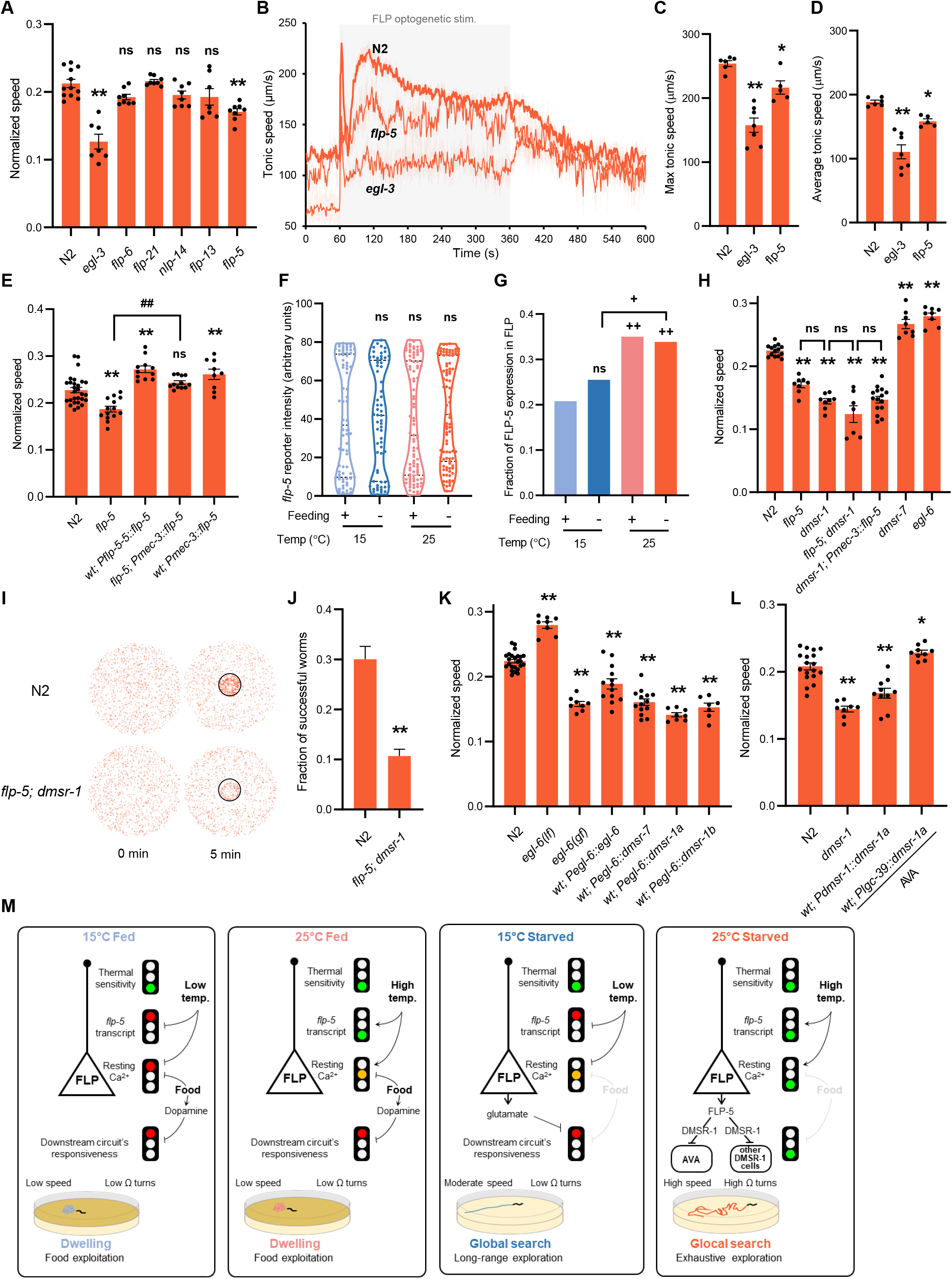
Multi-site ungating of tonic neuropeptide signaling in the FLP pathway controls speed elevation during glocal search. Impact of neuropeptide-affecting mutations on the speed of starved animals at 25°C (**A**) and after a tonic optogenetic FLP stimulation reported as in Fig. 5 (**B, C, D**). Impact of *flp-5* mutation, over-expression, and rescue/over-expression with *Pmec-3::flp-5* transgene expressed in FLP (**E**) and of mutations affecting FLP-5 receptors (**H**) on the speed of starved animals at 25°C. *flp-5* transcriptional reporter analysis reported as in Fig. 4 (**F-G**). Food drop assays reported as in Fig. 2 showing reduced food-reaching performances in *flp-5; dmsr-1* (**I, J**). Impact of gain-of-function (gf) and loss-of-function (lf) mutations in *egl-6*, as well as FLP-5 receptor over-expression in *egl-6-*expressing cells, revealing that EGL-6, DMSR-1a, DMSR-1b and DMSR-7 have a similar inhibitory effect on the speed of starved animals at 25°C (**K**). Opposite impact of broad DMSR-1 overexpression in *dmsr-1-*expressing cells or AVA-specific overexpression, respectively, on the speed of starved animals at 25°C (**L**). *, *p*<.05 and **, *p*<.01 versus 15°C Fed condition or respective N2 control; # *p*<.05 and ##, *p*<.01 versus the indicated condition by Bonferroni posthoc tests. +, *p*<0.05 and ++, *p*<.01 versus 15°C Fed or indicated condition by Fisher’s exact test for contingency comparisons. ns, not significant. Model for food and temperature-dependent control of exploitation/exploration strategies via multi-site gating mechanisms (traffic lights, **M**). On food, animals dwell regardless of temperature with one or more gates closed in the FLP-pathway (red lights). During global search after starvation at 15°C, only parts of the gates are lifted (green lights), still preventing the up-regulation of omega turn and of speed, the latter being also hindered by FLP glutamate signaling. After starvation at 25°C, all gates are lifted and FLP tonically signals via FLP-5/DMSR-1 to elevate speed and omega turns and orchestrate glocal search.

Both speed and omega turn defects were rescued by a *[Pmec-3::flp-5]* transgene, driving *flp-5* expression in FLP (Fig. 7E and S7B). Moreover, overexpression of *flp-5* under its native promoter or in FLP using the *mec-3* promoter caused further increase in speed and omega turn frequency compared to wild type (Fig. 7E, and S7B). While our data suggest several neuropeptides plays a role and do not rule out additional sources for FLP-5, they indicate that FLP-originating FLP-5 represents a relevant signal promoting the transition to glocal search.

### *flp-5* transcription is controlled by temperature in FLP

Since FLP-5 is a speed promoting signal from FLP, we next asked if *flp-5* transcription depends on temperature and/or feeding status. When imaging a *[Pflp-5::mNeonGreen]* transcriptional reporter line in FLP, we observed a bimodal distribution of signal intensity among cells with detectable expression that was similar in all conditions (Fig. 7F), but the fraction of animals with detectable *flp-5* reporter expression was significantly higher at 25°C compared to 15°C regardless of feeding status (Fig. 7G).

Considering that *flp-5* overexpression with *mec-3* promoter was sufficient to up-regulate speed and omega turns (Fig. 7E and S7B), we propose that reduced *flp-5* transcription in FLP at 15°C may be one of the gating mechanisms preventing the transition to glocal search at this temperature.

### FLP-5/DMSR-1 signaling is essential for glocal search

*In vitro* ligand screen data indicate that FLP-5 is a potent ligand of EGL-6 and DMSR-7 and activates DMSR-1 with lower potency (Beets et al., in prep). To identify functional FLP-5 receptors *in vivo*, we tested *egl-6, dmsr-7* and *dmsr-1* mutants. Speed and omega turn frequency was increased in *egl-6* and *dmsr-7* loss-of function mutants, which is opposite to the effect in *flp-5* mutant (Fig. 7H and S7C), suggesting that EGL-6 and DMSR-7 are relevant for these phenotypes, but are unlikely to directly mediate the glocal search-promoting effects of FLP-5. In contrast, *dmsr-1* loss-of-function mutants displayed decrease speed and omega turns frequency. *flp-5; dmsr-1* double mutants did not significantly differ from *dmsr-1* single mutants. Additionally, the speed and omega turn frequency increase caused by *flp-5* over-expression in FLP was blocked in a *dmsr-1* mutant background (Fig. 7H). We also confirmed the importance of FLP-5/DMSR-1 signaling for glocal search with our food drop assay, and showed that, as expected based on the marked impact on both speed and omega turn, the double *flp-5;dmsr-1* mutants performed very poorly (Fig. 7I-J). Taken together, these results are compatible with a model in which FLP-5 neuropeptide functions through DMSR-1 receptor to shift speed and omega turn frequency in order to support an efficient glocal search strategy.

### DMSR-1 and -7 function as inhibitory receptors *in vivo*

Next, we wanted to address if FLP-5 receptors work as excitatory or inhibitory receptors. EGL-6 is a well-established inhibitory receptor (Ringstad and Horvitz, 2008) and recent studies suggests an inhibitory role for DMSR-7 (Marquina-Solis et al., 2022) and DMSR-1 (Iannacone et al., 2017) as well. However, the two isoforms encoded by *dmsr-1* vary at their C-termini and could potentially activate different G-protein subtypes, calling for further analyses. We designed experiments to compare the effect of DMSR-1a/b and DMSR-7 to that of EGL-6. *egl-6(n4537)* loss-of-function mutation increased, whereas *egl-6(n592)* gain-of-function mutation decreased speed and omega turn frequency after starvation at 25°C (Fig. 7K, S7D). Overexpression of *egl-6* under its own promoter was also sufficient to down-regulate speed and omega turns. These results indicate that overexpressing an inhibitory receptor in *egl-6*-expressing cells negatively impacts speed and omega turn, hence providing a way to test if other receptors have the same inhibitory effect. The over-expression of DMSR-1a, DMSR-1b, as well as DMSR-7 all produced an inhibitory impact similar to EGL-6 over-expression (Fig. 7K and S7D). Therefore, the FLP-5 receptors identified *in vitro* (EGL-6, DMSR-1 and DMSR-7) display inhibitory activity *in vivo* when over-expressed in the circuit relevant for glocal search control.

### DMSR-1 activity in AVA command interneuron promotes speed elevation

Our next goal was to determine the locus of action of DMSR-1 receptor in controlling the speed elevation during glocal search. DMSR-1 is broadly expressed in the nervous system (Taylor et al., 2021), including in forward locomotion-promoting neurons (like AVB, RIB and DVA), as well as backward locomotion-promoting neurons (like AVA, whose activation was also shown to reduce animal speed (Schmitt et al., 2012)). Based on these observations, we made two predictions. First, since DMSR-1 is expressed in neurons with antagonistic impact on speed, then the FLP-5 signal relevant for the speed increase is unlikely to have a broad impact that generally affects all DMSR-1-expressing neurons. Consistent with this view, overexpressing DMSR-1a or b with its endogenous promoter failed to up-regulate speed and actually reduced it, suggesting that FLP-5 must act via a narrower subset of target cells in order to elevate speed. Second, since DMSR-1 is a low-affinity receptor and is inhibitory, its relevant place of action downstream of FLP-5 is likely to be speed-inhibiting neurons that are post-synaptic to FLP. AVA command interneuron was our top candidate and we found that AVA-specific overexpression of DMSR-1 primarily in AVA with the *lgc-39* promoter was indeed sufficient to increase speed (Fig. 7L). Taken together, these results elucidate an important signaling mechanism participating in the orchestration of the behavioral transition to glocal search, which involves the FLP-5/DMSR-1-dependent regulation of AVA by FLP.

## DISCUSSION

The ability to engage in persistent behavioral states and to switch between different states in response to external and internal multimodal cues is an essential animal capability to improve growth and survival via the execution of context-dependent behavioral strategies. Owing to the complexity of the signaling involved and of the behavioral responses themselves, we still know very little about the neural and molecular mechanisms that orchestrate coherent and long-lasting multidimensional changes, which permit to implement these behavioral strategies. Here, we systematically decode how food availability is integrated with past and recent thermosensory experience and show how these factors interact, leading the animal to select between different strategies. We confirmed previously described persistent behavioral states (dwelling and global search), highlighted novel exploitation and exploration states (scanning and glocal search, respectively) and illustrate their potential ecological benefits. Our dissection of the neural and molecular pathways involved provides two separate examples in which food and/or temperature signals converge to lift gates at multiple sites. This allows for tonic neuropeptide signaling by specific sensory neurons (AFD and FLP, respectively) to be only engaged when several conditions are met. The system dynamics might be tuned to cause animals to ignore short-lasting stimuli when committing to persistent behavioral states. But quite remarkably, the system still allows for a parallel control of short-term behavioral responses via phasic signaling by these same sensory neurons. The similarities between the two examples in AFD and FLP, respectively, suggest a potentially general regulatory logic through which sensory pathways integrate multimodal context to control behavioral responses over different timescales.

### Multimodal context-dependent exploitation/exploration strategies

Like in previous studies (Gallagher et al., 2013; Martineau et al., 2020), our results suggest that behavioral states under different contexts can be represented as a unique code of *C. elegans* locomotion and posture. On food, worms are in a dwelling state (Flavell et al., 2013; Fujiwara et al., 2002; Shtonda and Avery, 2006), regardless of the growth temperature. Thus, responses to different long-term temperature history are gated on food. Over a shorter timeframe, however, worms transiently enhanced dwelling state in response to cooling, while they persistently transition to a scanning state in response to warming. The asymmetry in the nature and duration of the responses to cooling and warming, respectively, reveals differential prioritization of environmental cues and specific behavioral strategies affecting the exploitation-exploration trade-off when adapting to a changing environment.

The scanning state during which animals produce more frequent bouts of slow forward and backward locomotion is different from the roaming behavior in which animals move faster and down-regulate reversals (Flavell et al., 2020). Whereas transition to scanning is less drastic than the transition to off-food search modes, animals in the scanning state sample their local environment more exhaustively than dwelling animals and, when confronted to a thermal gradient, produce more efficient thermotaxis. We speculate that short motion bouts during scanning may improve gradient detection, because motion-evoked thermal changes are the primary inputs through which worms decode their thermal environment (Clark et al., 2007). In previous studies, animal fitness was reduced when exposed to cycling temperature regimes, demonstrating that physiological adaption to new temperatures is not costless (Cedergreen et al., 2016). With little energy invested in locomotion and without disrupting local food exploitation, the scanning state could facilitate the detection of thermal gradients and help reach previous growth temperature in a slow-developing thermotactic drift that will limit the need for long-term physiological adaptations.

After prolonged starvation, current temperature influenced the persistent transition to previously known global search at 15 or to a new glocal search state at 25°C. During global search, animals increase forward movement at an intermediate speed and suppress turns, resulting in a long-range navigation with straight trajectories to locate food. During glocal search, concerted up-regulation of speed and omega turns results in a specific exploration strategy, where animals exhaustively sample local environment by making frequent turns while still performing long-range navigation due to marked speed increase. Glocal search enables finding food faster than global search, but is energetically expensive. We speculate that worm physiology and metabolism at 25°C puts worms under a higher time pressure, justifying the allocation of more resources for exploration.

As a whole, our study (i) expands the repertoire of described exploitation/exploration strategies in *C. elegans*, (ii) confirms the importance and efficacy of specific strategies, (iii) emphasizes the need for context-dependent transition between them, and (iv) expands our understanding of their multidimensional nature by comprehensively characterizing their underlying behavioral codes.

### Multisite gating in tonic sensory pathways integrates multimodal context

Although our data also suggest that additional pathways are involved, it is quite remarkable that the food and thermal contexts are largely encoded within AFD and FLP-dependent pathways. These mediate the transitions to scanning and to glocal search, respectively, and control both fast- and progressively-developing responses. Tonic signaling in these pathways, via specific neuropeptides, seems to have an instructive role and to be sufficient to trigger changes in many behavioral parameters. It is therefore important that robust gating mechanisms prevent their action under conditions where these strategies should not be engaged.

The transition to scanning requires AFD, which signals via FLP-6 neuropeptide. In scanning animals, AFD thermal responsiveness to short thermal stimuli is not affected, but AFD is in a particular state combining (i) sustained tonic activity (with high resting calcium levels) and (ii) increased *flp-6* expression. During dwelling at constant temperature, neither increased AFD resting calcium (at 15°C), nor increased *flp-6* transcript levels (at 25°C) seem sufficient to trigger scanning entry. Only after a recent warming, does AFD shows synergistic increases in steady state calcium and *flp-6* transcript levels (green lights in Fig. 4L). Whereas our data do not rule out additional modulation mechanisms downstream of AFD, they are consistent with a model in which multiple intracellular gates located at the sensory neuron level need to be lifted in order for the animal to shift its behavioral state (Fig. 4L and S8).

The transition to glocal search is promoted by FLP, via FLP-5 and most likely also via additional unidentified neuropeptides. The FLP-pathway integrates temperature and food signals to promote the transition to glocal search only at high temperature and in the absence of food. Like for AFD, the FLP thermal responsiveness to short stimuli was not different between behavioral states, indicating that the distinction between states is encoded downstream of the FLP thermo-detection process. Glocal search was associated with (i) a very high resting calcium level in FLP, (ii) elevated *flp-5* transcription and (iii) increased responsiveness of the circuit downstream of FLP activation (Fig. 7M). Given the likely instructive role that tonic FLP-5 signaling from FLP has in the transition to glocal search, each of these three processes might represent a potential gate able to prevent this transition (Fig. 7M). Indeed, in dwelling animals at 15°C on food, all three gates are closed with low baseline FLP calcium, low *flp-5* transcript and low responsiveness of components downstream of FLP. In animals performing global search at 15°C off-food, we only observed a partial increase in FLP steady-state calcium levels, while the rest of the gates were unchanged compared to dwelling animals on food at 15°C. Furthermore, glutamate signaling from FLP contributes to prevent speed increase. Several neurons (ASI, ASK, AIA, AVK) (Gray et al., 2005; Lopez-Cruz et al., 2019; Oranth et al., 2018) have been implicated in transition to global search and probably FLP signals information of low temperature in this context. In dwelling animals at 25°C on food, we observed a partial increase in FLP resting calcium levels and an increase in *flp-5* transcript, but dopamine signaling reduced the responsiveness of components downstream of FLP, which might be the most important gate in this context in order to block the high temperature signal. Ultimately, the glocal search mode is associated with a unique state in the FLP pathway where food absence and recent temperature signals converge to lift all the cellular gates in order to produce context-specific persistent behavioral transitions (Fig. 7M).

Taken together, the two examples of AFD- and FLP-dependent behavioral transitions described here, suggest a generally applicable regulatory logic, where multisite gates, qualitatively and quantitatively affecting tonic signaling and its reading in sensory circuits, function as “relay boards” for temporal and multimodal integration of signals. The system will thus enable behavioral transitions to occur only in response to coherent and sustained changes in the environment.

### A distributed network of inhibitory GPCR(s) orchestrates behavioral transitions

The progressive increase in complexity throughout our study limited our ability to dissect all the downstream molecular and cellular components of the entire behavioral code that is modulated during different behavioral state transitions. However, the sequential implementation of a reductionist analytical approach allowed us to test directed hypotheses and identify functional receptors for FLP-6 and FLP-5 as well as one relevant cellular locus of action for the latter, hence providing a partial, yet suggestive picture of the downstream effector functional logic.

Tonic FLP-6 signaling from AFD was previously shown to mediate the temperature effect on lifespan via unknow receptors (Chen et al., 2016). Here we show that FLP-6 might promote the transition to scanning, through at least three inhibitory GPCRs broadly expressed in the nervous system: EGL-6, known to regulate egg-laying (Ringstad and Horvitz, 2008), DMSR-1, known to regulate stress-induced sleep (Iannacone et al., 2017), and DMSR-7, recently proposed to control sickness behavior (Marquina-Solis et al., 2022). Our data suggest that FLP-6 mediates a progressive increase in backward frequency through EGL-6 and a fast decrease in tail bending through DMSR-7, DMSR-1, and EGL-6. Therefore, different FLP-6 receptors contribute to regulate different behavioral parameters over different timescales to ensure transition and maintenance of the scanning state.

Our study also sheds light on some downstream mediators of the FLP pathways. During transition to glocal search, FLP-5 functions through inhibitory DMSR-1 by inhibiting AVA and unidentified neurons to increase speed and omega turns, respectively. Interestingly, speed up-regulation entails the inhibition of an inhibitory pathway, suggesting that the animal default state under favorable conditions would be a metabolically inexpensive low arousal state. Moreover, fine-tuning the neural circuit activity by modulating the inhibition of the key command interneuron AVA (Kato et al., 2015; Liu et al., 2020) can enable the transition to a metabolically expensive but adaptive persistent behavioral state.

In conclusion, our study suggests that *C. elegans* continuously weights the sensory valence of temperature in distinct tonic thermosensory neurons, based on past experience, as well as internal and external contexts. In the thermosensory circuit, multi-site cellular and molecular gatings act in concert to generate physiologically adaptive behavioral states by balancing exploration-exploitation strategies.

Moreover, only in specific contexts does the GPCR(s) dependent fine-tuning of excitation-inhibition balance mediate persistent behavioral state transitions. Given the prevalence of tonic sensory signaling, the widespread expression of GPCRs and their emerging functions in brain state pathologies (i.e., anxiety, depression, schizophrenia) (Belzeaux et al., 2020; Magalhaes et al., 2010; Matsumoto et al., 2008), we propose that similar multi-site gating-dependent integration mechanisms converging on GPCRs might mediate persistent behavioral state transitions in response to a variety of internal and external cues and in additional species.

## ACKNOWLEDGMENTS

The authors thank Lisa Schild and Laurence Bulliard for technical support; Gabriella Saro for help with calcium imaging; Andrei-Stefan Lia for help in setting-up the optogenetic device; Andre Brown for help with the Tierpsy tracker; Yaïleen Bonvin, Wayne Davis, Miriam Goodman, Marc Hammarlund, Domenica Ippolito, Andreina Rueda, Bill Schafer and Piali Sengupta for the gift of strains and plasmids; Elise-Dan-Glauser, Simon Sprecher, and Arjit Kant Gupta for comment on the manuscript. Some strains were provided by the CGC, which is funded by NIH Office of Research Infrastructure Programs (P40 OD010440). The study was supported by the Swiss National Science Foundation (IZCNZ0_174703/SBFI_C16.0013, BSSGI0_155764 and PP00P3_150681 to D.A.G) and by the Research Foundation – Flanders (FWO G093419N to I.B.).

## AUTHOR CONTRIBUTIONS

Study conceptualization and experiment design (ST, DAG), methodology development (ST, DAG), execution of all experiments (ST) except for *in vitro* neuropeptide GPCR interaction analyses (IB), data analysis and interpretation (ST, DAG), article writing (ST, DAG), supervision (DAG), funding acquisition (DAG, IB).

## METHODS

### *C elegans* growth, maintenance and synchronization

*C. elegans* strains were maintained according to standard techniques on nematode growth medium (NGM) agar plates seeded with OP50. Animal synchronization was made by treating gravid adult with standard hypochlorite-based procedure. Strain List section below includes a list of strains used in the present study.

### Behavior analyses

An overview of the device and pipeline for behavioral recording and analyses is presented in Fig. S1A and B, and detailed below.

### Behavioral recordings

#### Video recordings

Behavior of animal populations crawling on solid medium plates was recorded in a custom-made temperature, vibration and illumination-controlled platform (Figure S1A). High-resolution (2448×2048 pixels), 3-min movies (∼50 animals/movie) were recorded using DMK33UX250 camera (The Imaging Source), at 10 frames per second as a snapshot of the behavioral state in given condition using IC Capture software (The Imaging Source) and saved as .*avi* files. All the behavioral recordings across conditions and genotypes were performed in young adult animals (∼800 µm in length). Each condition and genotype were recorded on at least 3 separate days with multiple replicates each day. All the behavioral experiments were performed in standard 6-cm NGM petri dishes (with or without *E. coli* food lawn) in a lid-on setting unless otherwise mentioned.

#### Steady-state behavior

Animals were grown and recorded without changing the temperature throughout life for steady-state/baseline behavior at 15°C or 25°C, respectively. For detailed behavioral state characterization young-adult wild type animals were recorded at 15°C from 93-99hr, and at 25°C from 49-55hr for 6 hours. For mutants, steady state baseline behavior was recorded once in young adult animals.

#### Temperature shift experiments

For temperature shift experiments, animals were moved to the recording device pre-set at target temperature just before the recording. Target temperature of 15°C or 25°C was achieved within 45 minutes after temperature shift. Duration of transition was counted from the time of plate shift. Wild-type animals for detailed behavioral state characterization were recorded every hour for 6 hours. Mutants were only recorded after 6hr of temperature shift to focus on steady state behavioral states.

#### Starvation experiments

Fed animals were washed 3 times in the 1.5mL collection tube with distilled water (pre-equilibrated at growth temperature of worms) to remove OP50. During washes worms were left to settle to the bottom of the plates by gravity. About 50-80 animals were then plated on NGM plates with a drop of water and left to airdry for 5 minutes. Total duration of wash and drying was ∼10 min per plate. Duration of starvation was considered from the time of start of the wash. Wild-type animals for detailed behavioral state characterization were recorded every hour for 6 hours. Mutants were only recorded after 6hr of starvation shift to focus on steady state behavioral states.

#### Preparing transgenic animals for behavioral recordings

Transgenic animals were picked 24 h before recordings and placed to equilibrate in the recording device.

### Behavioral parameter extraction from behavioral movies

Recorded movie files were analysed using the Tierpsy tracker v1.4 for detailed quantification of locomotion and posture (Javer et al., 2018a). The output feature file contains average value for ∼700 behavioral parameters for all tracked worms. Previous work recommended using a smaller parameter subset with 256 Tierpsy features for balancing power and interpretability (Javer et al., 2018b). We reasoned that this list of 256 parameters is suitable for the studies where high-power is required to capture all the possible phenotypic difference (e.g. genetic or drug screens). However, this total number of is still relatively large to handle in a study performing multiple subsequent mechanistic dissections and still contains many parameters which are either (i) difficult to interpret, (ii) inherently noisy, or (ii) still redundant with others. We used an alternative approach to decrease the list of Tierpsy v1.4 parameters (∼700) using the following reasoning:

1. Include all the parameters accounting for major postural and locomotion variations and likely to have a neural basis and an interpretable ecological impact (bending of posture at different body parts, eigen projections, speed, omega turns, reversals, forward and pause locomotion)
2. Remove redundancy wherever possible (only used abs value and excluded positive and negative value of the same parameter)
3. Remove noisy parameters with empty values for most datapoints (as upsilon turn frequency).
4. Keep sufficient parameters to grasp complete information about a behavioral cluster (backward frequency, backward time, inter backward time, backward time ratio)
5. Further remove redundant parameters of a behavioral cluster likely to have same mechanistic basis (removed backward distance, inter backward distance, backward distance ratio based on the logic that backward time and distance parameters might have same underlying mechanism)
6. Only keep information of a behavioral parameter for individual body part where required (e.g., all the bending parameters kept for individual body part) else only information of midbody is used (e.g., only keep midbody speed but not tail speed).
7. Length normalized speed was introduced later in the study to control for growth/age dependent variations across conditions and genotypes.

Applying the reasoning, we focused on a set of 47 (17 postural, and 30 motion, as described in Fig. S1C) readily interpretable behavioral parameters, which we reasoned best explain all the changes, likely to have a directly addressable mechanistic basis and can represent behavioral states of the animals.

### PCA and Hierarchical clustering

We performed a global Principal component analysis (PCA) on raw average values of 47 behavioral parameters examined across all the wild type conditions in our study. PCA on motion only was performed on 30 parameters and on posture only was performed on 17 postural parameters (Figure S1C). Phenotypic clustering was performed on *z*-score normalized values of each parameter across all conditions for all the behavioral parameters. PCA and hierarchical clustering was performed using Clustvis webtool (Metsalu and Vilo, 2015). This integrated quantitative comparison across all the conditions allowed us to map behavioral states in a common multidimensional space.

### Optogenetics

Optogenetics experiments in temperature-controlled environment, were performed using a custom-made plug-in device attached to our high-content behavioral recording platform (Fig. S1A).

Homogeneous blue light stimuli were delivered though a ring of blue LED (460 ± 10 nm). Power was adjusted with an optocoupler-based system and the delivered light intensity (15 W/m^2^) was determined with a portable spectrophotometer (USB4000, OceanOptics). Animals were grown on all-trans-retinal (ATR) plates as previously described (Schild and Glauser, 2015). We previously published control experiment in the absence of ATR showing no behavioral response at this light intensity (Saro et al., 2020). Animals were recorded for 600 s (60 s baseline, 300 s light-stimulation, 240 s recovery).

Maximum and average tonic speed elevation were calculated only during for the time span between *t=*90 and 360 s in order to exclude the first phase of reversal bursts observed at the onset of FLP stimulation. The speed data for optogenetic experiments were obtained as Tierpsy timeseries feature output ‘abs midbody speed’ and represents absolute average speed (in forward or backward locomotion) of all the animals at each timepoint.

### Locomotion trajectory analysis and simulation

Real locomotion trajectories (1 min) were calculated based on x-y coordinate data of 35 randomly sampled animals in each behavioral state. Trajectory Monte-Carlo simulations were performed with previously published custom-made Excel sheets (Schild and Glauser, 2013), emulating an isothermal environment by simply setting the spatial thermal gradient slope to 0. Distance represents summed path travelled over 1 min and displacement represents the length of a linear path between starting and end point of the trajectory.

### On-food thermotaxis assay

The on-food thermotaxis assay is a modified version of previously described thermotaxis assays on linear gradient usually carried out on food-free plates, with freshly food-deprived animals and lasting < 1 h (Goodman et al., 2014). Here, experiments were performed on food, without worm transfer procedure and lasted longer (up to 3 h). Animals were transferred to experimental NGM plates completely and evenly covered with OP50 24 h before recordings. Animals maintained at mentioned isothermal conditions were gently transferred to a stable linear thermal gradient (0.5°C/cm) created on an aluminium plate with a previously described custom-made thermalized system (Schild and Glauser, 2013). Following the plate transfer, the thermal gradient was established within 10 min. Movement was captured once before and at multiple timepoints after transfer using DMK 21BU04 camara (The imaging source). The entire experiment was performed in a lid-off setting to monitor temperature in the gradient using an infrared thermometer. Coordinate data of worms were obtained by manually spotting in each image using ImageJ. On-food thermotaxis index was calculated as the mean of the population distribution normalized to the plate size (values ranging from 0 to 1). An index value of 0 would correspond to an extreme cryophilic bias with all the worms at the cool end of the plate; a value of 1 would correspond to an extreme thermophilic bias with all the worms at the warm end of the plate; a value, 0.5 corresponds to an even distribution on both side of the plate center (like at the start of each experiment).

### Food drop assay

Animals were starved at 15°C or 25°C on 6-cm NGM plates as described above. After 6 h of starvation a dense 20 µL drop of OP50 (OD600 after 1:100 dilution: 0.57A) was dropped at the center of the plate. Experiments were performed in lid-on settings and lid was only removed once to add the OP50 drop. Snapshots of worm distribution were taken before and after at desired timepoints. Coordinate data of worms was obtained by manually spotting worms in each image using ImageJ. Region of success was defined as a circular region of 1.6 cm diameter around the food drop. Animals already present at the region of success before dropping the food were discarded from the analysis. Fraction of successful food search was calculated using the following formula:

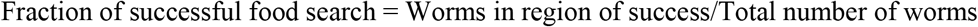

### Calcium imaging

Calcium imaging experiments in AFD and FLP were performed in a temperature-controlled environment and analysed as previously described (Ippolito et al., 2021; Saro et al., 2020). YFP/CFP ratio of a YC2.3 cameleon indicator was used to compare resting calcium levels across conditions, as well as relative changes in response to short-lasting stimuli. When the impact of starvation and thermal shift were examined, recordings were made 6 h after the condition changes.

### Microscopy and reporter expression analysis

Images to quantify *flp-5* and *flp-6* reporter expression in FLP and AFD respectively, were acquired in a Zeiss Axioplan2 fluorescence microscope, with a 40× (air, NA = 0.95) objective and constant illumination parameters.

### Statistical tests

D’Agostino & Pearson test (p<0.01) was used to test normality of distributions. Comparisons giving significant effects (p<0.05) with ANOVAs were followed by Bonferroni posthoc tests. Dunn’s test was used as non-parametric test whenever the normality assumption criterion was not fulfilled. In every figure, statistical significance is represented as ns, not significant, *, *p*<0.05 and **, *p*<0.01 for comparison with N2 15 fed or respective N2 controls and #, *p*<0.05 and ##, *p*<0.01 for indicated comparison. The χ2 and Fisher’s exact tests were performed for discrete event contingency comparisons and statistical significance is represented as ns, not significant, +, *p*<0.05 and ++, *p*<0.01. Bimodal distribution was represented as violin plot and the rest of the data is represented as bar graph with individual datapoints overlaid.

### Transgene construction and transgenesis

DNA prepared with a GenElute HP Plasmid miniprep kit (Sigma) was microinjected in the gonad to generate transgenic lines according to a standard protocol (Evans, 2006). We used a [*unc-122p*::*GFP*] (gift from Piali Sengupta; Addgene plasmid # 8937 (Miyabayashi et al., 1999)) co-injection marker to identify transgenic animals. The concentration of co-marker and for expression plasmids in the injection mixes are indicated in the strain list.

### Promoter plasmids (MultiSiteGateway slot 1)

Entry plasmids containing specific promoters were constructed by PCR from N2 genomic DNA with primers flanked with attB4 and attB1r recombination sites and cloned into pDONR-P4-P1R vector (Invitrogen) by BP recombination. Primer sequences were the following:

dg507 [slot1 Entry gcy-8p]

attB4gcy-8_F: ggggacaactttgtatagaaaagttgATAGCAAAGGGCGTCGATTATCT

attB1rgcy-8_R: ggggactgcttttttgtacaaacttgTTTGATGTGGAAAAGGTAGAATCGAA

dg867 [slot1 Entry egl-6p]

attB4egl-6_F: ggggacaactttgtatagaaaagttgATTTTCCAGAGAGAACAGAGTCC

attB1regl-6_R: ggggactgcttttttgtacaaacttgTTGCTGAAAAGCTGTCATTGTG

dg868 [slot1 Entry flp-5p]

attB4flp-5_F: ggggacaactttgtatagaaaagttgATCGAATTTGTCGCCGATCTGTTACA

attB1rflp-5_R: ggggactgcttttttgtacaaacttgTAGTTGCGAGGAATGACTGTTTCG

dg865 [slot1 Entry dmsr-1p]

attB4dmsr-1_F: ggggacaactttgtatagaaaagttgATCAGACGTCGTTGTGGAAGTAG

attB1rdmsr-1_R: ggggactgcttttttgtacaaacttgTTTTGTTTGCTGTTCCTCTGTTC

dg1015 [slot1 Entry lgc-39p]

attB4lgc-39_F: ggggacaactttgtatagaaaagttgATCGCTATCATCGTCTCCAAATCG

attB1rlgc-39_R: ggggactgcttttttgtacaaacttgTCGATGATTCACATCAGGGATGC

The generation of dg68 [slot1 Entry mec-3p] was previously described (Schild et al., 2014).

### Coding sequence plasmids (MultiSiteGateway slot 2)

Entry plasmids containing specific coding DNA sequences (cds) or genomic sequence (gs) were constructed by PCR from N2 cDNA or N2 genomic DNA with primers flanked with attB1 and attB2 recombination sites and cloned into pDONR_221 vector (Invitrogen) by BP recombination. Primer sequences were the following:

dg1014 [slot2 Entry flp-6cds]

attB1flp-6_F: ggggacaagtttgtacaaaaaagcaggctTAATGAACTCTCGTGGGTTGATTTTGA

attB2flp-6_R: ggggaccactttgtacaagaaagctgggtCTTATCGTCCGAATCTCATGTATGCT

dg872 [slot2 Entry egl-6gs]

attB1egl-6_F: ggggacaagtttgtacaaaaaagcaggctTAATGAATGACACACTGATCTGTACA

attB2egl-6_R: ggggaccactttgtacaagaaagctgggtCTTAAGACCCGACATATGAGCTTG

dg875 [slot2 Entry flp-5cds]

attB1flp-5_F: ggggacaagtttgtacaaaaaagcaggctTAATGAGCAGCCGAAGCACCAC

attB2flp-5_R: ggggaccactttgtacaagaaagctgggtCTTAGCCGAATCGGATGAATTTGGCT

dg871 [slot2 Entry dmsr-7cds]

attB1dmsr-7_F: ggggacaagtttgtacaaaaaagcaggctTAATGGAATGTCCGCACGATGC

attB2dmsr-7_R: ggggaccactttgtacaagaaagctgggtCTTAAAGTTGATGTTCTCTACTGCTG

dg869 [slot2 Entry dmsr-1Acds]

attB1dmsr-1a_F: ggggacaagtttgtacaaaaaagcaggctTAATGGAGTTTACCGAATGCAAAAC

attB2dmsr-1a_R: ggggaccactttgtacaagaaagctgggtCTCAAATGTTTTGAAAGTGTCCACG

dg870 [slot2 Entry dmsr-1Bcds]

attB1dmsr-1b_F: ggggacaagtttgtacaaaaaagcaggctTAATGGAGTTTACCGAATGCAAAAC

attB2dmsr-1b_R: ggggaccactttgtacaagaaagctgggtCTTATTTCCGTACTGTTTCTTCGTAC

dg88 [slot2 Entry TeTxcds] (aka pWD157) was a gift of Wayne Davis. The generation of dg651 [slot2 Entry egl-13NLS::wrmScarlet], of dg353 [slot2 Entry mNeongreen cds] and dg650 [slot2 Entry NLS_ceBFPcds] were described in (Marques et al., 2019), (Hostettler et al., 2017) and (Ippolito et al., 2021) respectively.

### 3’ UTR and tagging plasmids (Multi-site Gateway slot3)

mg277 [SL2::mCherry] was previously described (Schild et al., 2014). mg211 [EntrySlot3unc-54UTR] (aka pMH473) was a gift from Marc Hammarlund.

### Expression plasmids used for transgenesis

dg931 [gcy-8p::QF::unc-54UTR] was created through a LR recombination reaction between dg507, dg240, mg211, and pDEST-R4-P3.

dg899 [egl-6p::QF::unc-54UTR] was created through a LR recombination reaction between dg867, dg240, mg211, and pDEST-R4-P3.

dg243 [mec-3p::QF::unc-54UTR] was created through a LR recombination reaction between dg68, dg240, mg211, and pDEST-R4-P3.

dg898 [flp-5p::QF::unc-54UTR] was created through a LR recombination reaction between dg868, dg240, mg211, and pDEST-R4-P3.

dg900 [dmsr-1p::QF::unc-54UTR] was created through a LR recombination reaction between dg865, dg240, mg211, and pDEST-R4-P3.

dg1017 [lgc-39p::QF::unc-54UTR] was created through a LR recombination reaction between dg1015, dg240, mg211, and pDEST-R4-P3.

dg255 [QUASp::TeTxcds::SL2::mCherry] was created through a LR recombination reaction between dg229, dg88, mg277, and pDEST-R4-P3.

dg839 [QUASp::flp-6cds::SL2::mCherry] was created through a LR recombination reaction between dg229, dg827, mg277, and pDEST-R4-P3.

dg894 [QUASp::egl-6gs::SL2::mCherry] was created through a LR recombination reaction between dg229, dg872, mg277, and pDEST-R4-P3.

dg838 [QUASp::eat-4cds::SL2::mCherry] was created through a LR recombination reaction between dg229, dg713, mg277, and pDEST-R4-P3.

dg893 [QUASp::flp-5cds::SL2::mCherry] was created through a LR recombination reaction between dg229, dg875, mg277, and pDEST-R4-P3.

dg895 [QUASp::dmsr-7cds::SL2::mCherry] was created through a LR recombination reaction between dg229, dg871, mg277, and pDEST-R4-P3.

dg897 [QUASp::dmsr-1Acds::SL2::mCherry] was created through a LR recombination reaction between dg229, dg869, mg277, and pDEST-R4-P3.

dg896 [QUASp::dmsr-1Bcds::SL2::mCherry] was created through a LR recombination reaction between dg229, dg870, mg277, and pDEST-R4-P3.

dg373 [QUASp::mNeongreencds::unc-54UTR] was created through a LR recombination reaction between dg229, dg353, mg211, and pDEST-R4-P3.

dg996 [QUASp::wrmScarletcds::unc-54UTR] was created through a LR recombination reaction between dg229, dg353, mg211, and pDEST-R4-P3.

dg653 [mec-3p::NLS_CeBFPcds::unc-54UTR] was created through a LR recombination reaction between dg68, dg650, mg211, and pDEST-R4-P3.

### Strain List

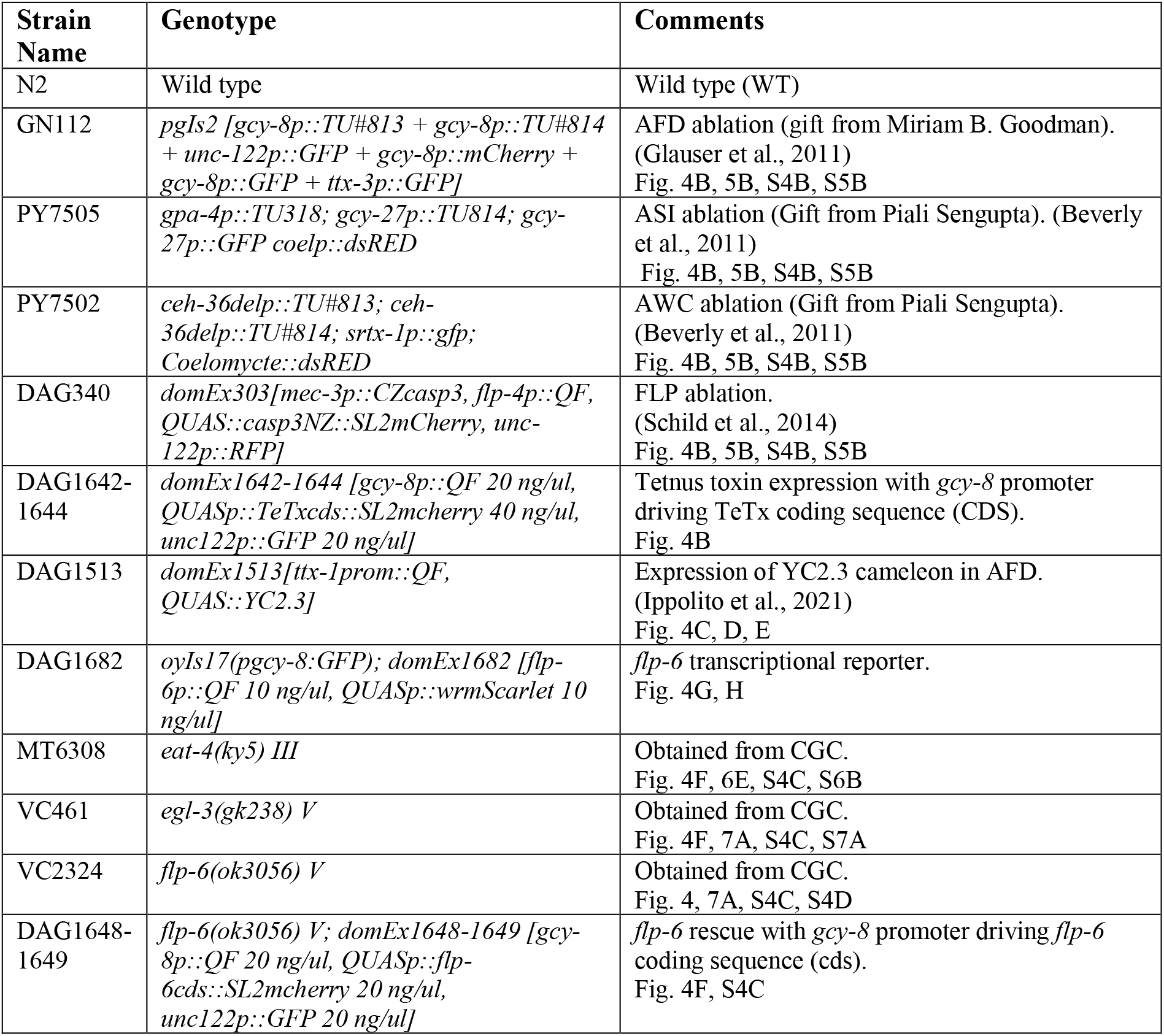
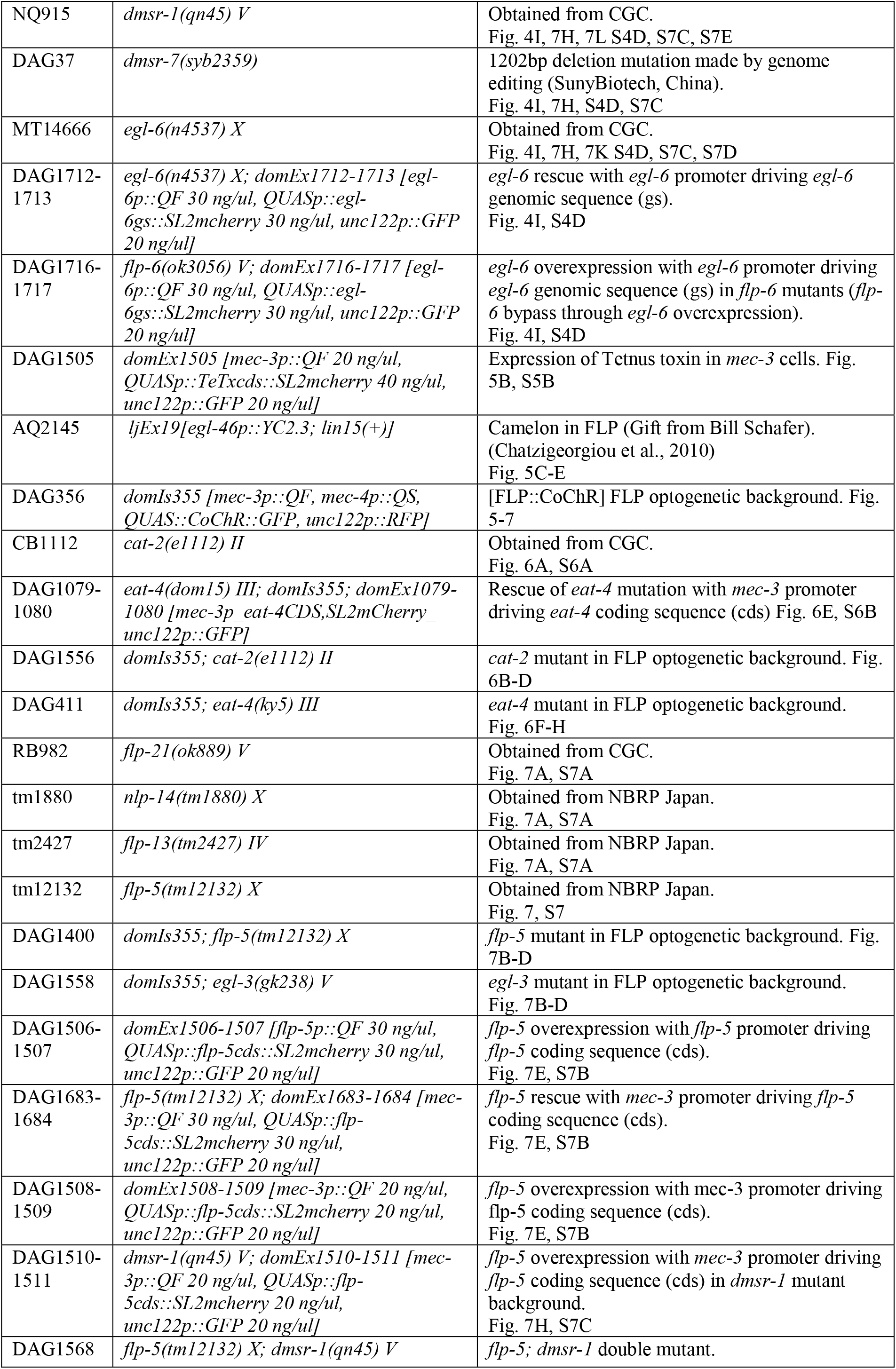
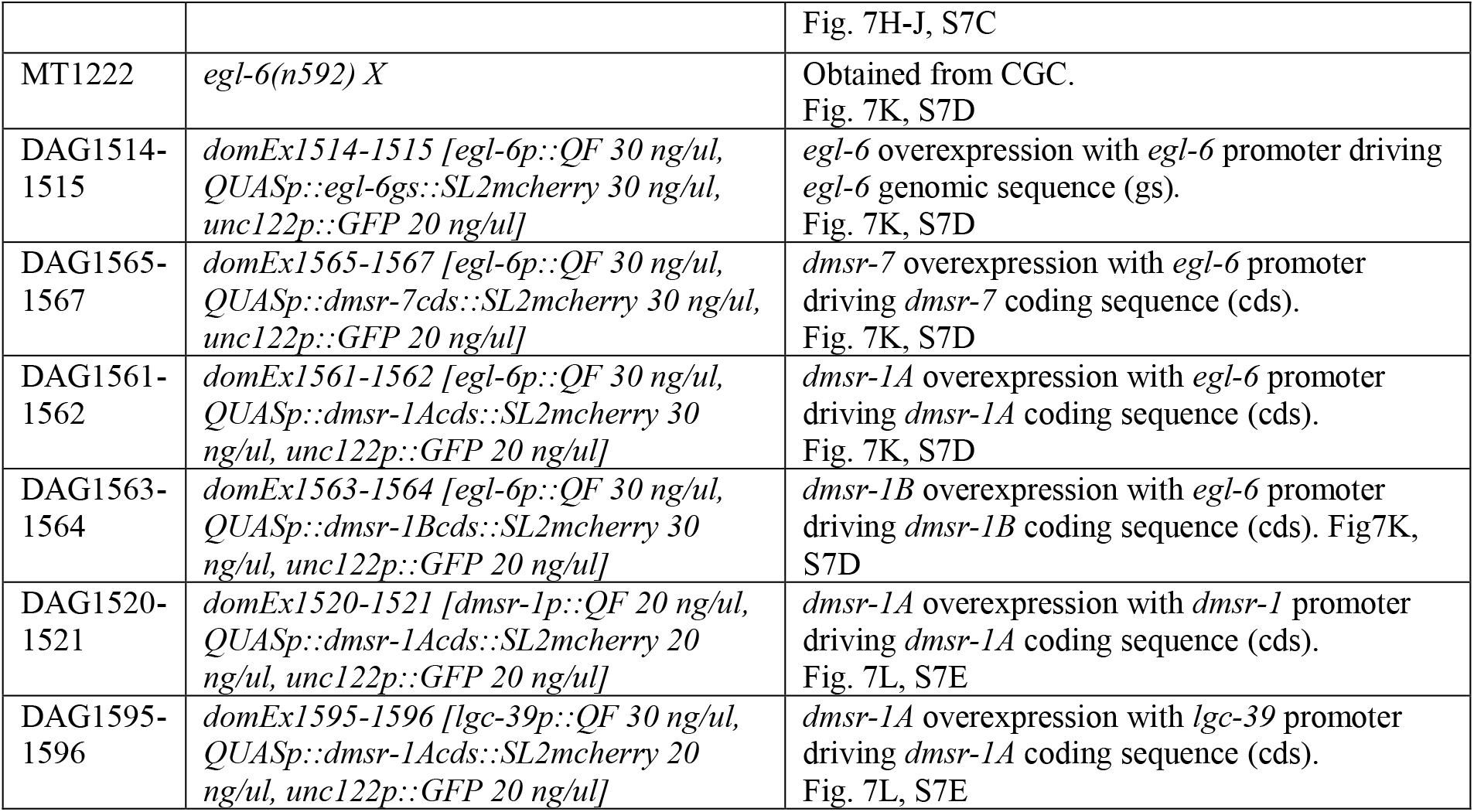

## REFERENCES

Andalman, A.S., Burns, V.M., Lovett-Barron, M., Broxton, M., Poole, B., Yang, S.J., Grosenick, L., Lerner, T.N., Chen, R., Benster, T., et al. (2019). Neuronal Dynamics Regulating Brain and Behavioral State Transitions. Cell 177, 970–985 e920.

Aoki, I., and Mori, I. (2015). Molecular biology of thermosensory transduction in C. elegans. Curr Opin Neurobiol 34, 117–124.

Bargmann, C.I. (2006). Chemosensation in C. elegans. WormBook, 1–29.

Belzeaux, R., Gorgievski, V., Fiori, L.M., Lopez, J.P., Grenier, J., Lin, R.X., Nagy, C., Ibrahim, E., Gascon, E., Courtet, P., et al. (2020). GPR56/ADGRG1 is associated with response to antidepressant treatment. Nature Communications 11.

Berridge, C.W., and Waterhouse, B.D. (2003). The locus coeruleus-noradrenergic system: modulation of behavioral state and state-dependent cognitive processes. Brain Res Brain Res Rev 42, 33–84.

Beverly, M., Anbil, S., and Sengupta, P. (2011). Degeneracy and neuromodulation among thermosensory neurons contribute to robust thermosensory behaviors in Caenorhabditis elegans. J Neurosci 31, 11718–11727.

Bhardwaj, A., Thapliyal, S., Dahiya, Y., and Babu, K. (2018). FLP-18 Functions through the G-Protein-Coupled Receptors NPR-1 and NPR-4 to Modulate Reversal Length in Caenorhabditis elegans. J Neurosci 38, 4641–4654.

Bhat, U.S., Shahi, N., Surendran, S., and Babu, K. (2021). Neuropeptides and Behaviors: How Small Peptides Regulate Nervous System Function and Behavioral Outputs. Front Mol Neurosci 14, 786471.

Bramham, C.R., and Srebro, B. (1989). Synaptic plasticity in the hippocampus is modulated by behavioral state. Brain Res 493, 74–86.

Bretscher, A.J., Busch, K.E., and de Bono, M. (2008). A carbon dioxide avoidance behavior is integrated with responses to ambient oxygen and food in Caenorhabditis elegans. Proc Natl Acad Sci U S A 105, 8044–8049.

Busch, K.E., Laurent, P., Soltesz, Z., Murphy, R.J., Faivre, O., Hedwig, B., Thomas, M., Smith, H.L., and de Bono, M. (2012). Tonic signaling from O(2) sensors sets neural circuit activity and behavioral state. Nat Neurosci 15, 581–591.

Cedergreen, N., Norhave, N.J., Svendsen, C., and Spurgeon, D.J. (2016). Variable Temperature Stress in the Nematode Caenorhabditis elegans (Maupas) and Its Implications for Sensitivity to an Additional Chemical Stressor. PLoS One 11, e0140277.

Cermak, N., Yu, S.K., Clark, R., Huang, Y.C., Baskoylu, S.N., and Flavell, S.W. (2020). Whole-organism behavioral profiling reveals a role for dopamine in state-dependent motor program coupling in C. elegans. Elife 9.

Chatzigeorgiou, M., Yoo, S., Watson, J.D., Lee, W.H., Spencer, W.C., Kindt, K.S., Hwang, S.W., Miller, D.M., 3rd, Treinin, M., Driscoll, M., et al. (2010). Specific roles for DEG/ENaC and TRP channels in touch and thermosensation in C. elegans nociceptors. Nat Neurosci 13, 861–868.

Chen, Y.C., Chen, H.J., Tseng, W.C., Hsu, J.M., Huang, T.T., Chen, C.H., and Pan, C.L. (2016). A C. elegans Thermosensory Circuit Regulates Longevity through crh-1/CREB-Dependent flp-6 Neuropeptide Signaling. Dev Cell 39, 209–223.

Churgin, M.A., McCloskey, R.J., Peters, E., and Fang-Yen, C. (2017). Antagonistic Serotonergic and Octopaminergic Neural Circuits Mediate Food-Dependent Locomotory Behavior in Caenorhabditis elegans. J Neurosci 37, 7811–7823.

Clark, D.A., Gabel, C.V., Gabel, H., and Samuel, A.D. (2007). Temporal activity patterns in thermosensory neurons of freely moving Caenorhabditis elegans encode spatial thermal gradients. J Neurosci 27, 6083–6090.

Dave, A.S., Yu, A.C., and Margoliash, D. (1998). Behavioral state modulation of auditory activity in a vocal motor system. Science 282, 2250–2254.

Devineni, A.V., and Scaplen, K.M. (2021). Neural Circuits Underlying Behavioral Flexibility: Insights From Drosophila. Front Behav Neurosci 15, 821680.

Evans, T. (2006). Transformation and microinjection (April 6, 2006), WormBook, ed. The C. elegans Research Community, WormBook, doi/10.1895/wormbook. 1.108. 1.

Flavell, S.W., Pokala, N., Macosko, E.Z., Albrecht, D.R., Larsch, J., and Bargmann, C.I. (2013). Serotonin and the neuropeptide PDF initiate and extend opposing behavioral states in C. elegans. Cell 154, 1023–1035.

Flavell, S.W., Raizen, D.M., and You, Y.J. (2020). Behavioral States. Genetics 216, 315–332.

Fu, X., Teboul, E., Weiss, G.L., Antonoudiou, P., Borkar, C.D., Fadok, J.P., Maguire, J., and Tasker, J.G. (2022). Gq neuromodulation of BLA parvalbumin interneurons induces burst firing and mediates fear-associated network and behavioral state transition in mice. Nat Commun 13, 1290.

Fujiwara, M., Sengupta, P., and McIntire, S.L. (2002). Regulation of body size and behavioral state of elegans by sensory perception and the EGL-4 cGMP-dependent protein kinase. Neuron 36, 1091–1102.

Gallagher, T., Bjorness, T., Greene, R., You, Y.J., and Avery, L. (2013). The geometry of locomotive behavioral states in C. elegans. PLoS One 8, e59865.

Ghosh, D.D., Lee, D., Jin, X., Horvitz, H.R., and Nitabach, M.N. (2021). C. elegans discriminates colors to guide foraging. Science 371, 1059–1063.

Gibson, W.T., Gonzalez, C.R., Fernandez, C., Ramasamy, L., Tabachnik, T., Du, R.R., Felsen, P.D., Maire, M.R., Perona, P., and Anderson, D.J. (2015). Behavioral responses to a repetitive visual threat stimulus express a persistent state of defensive arousal in Drosophila. Curr Biol 25, 1401–1415.

Glauser, D.A. (2022). Temperature sensing and context-dependent thermal behavior in nematodes. Curr Opin Neurobiol 73, 102525.

Glauser, D.A., Chen, W.C., Agin, R., Macinnis, B.L., Hellman, A.B., Garrity, P.A., Tan, M.W., and Goodman, M.B. (2011). Heat avoidance is regulated by transient receptor potential (TRP) channels and a neuropeptide signaling pathway in Caenorhabditis elegans. Genetics 188, 91–103.

Goodman, M.B., Klein, M., Lasse, S., Luo, L., Mori, I., Samuel, A., Sengupta, P., and Wang, D. (2014). Thermotaxis navigation behavior. WormBook, 1–10.

Goodman, M.B., and Sengupta, P. (2019). How Caenorhabditis elegans Senses Mechanical Stress, Temperature, and Other Physical Stimuli. Genetics 212, 25–51.

Gray, J.M., Hill, J.J., and Bargmann, C.I. (2005). A circuit for navigation in Caenorhabditis elegans. Proc Natl Acad Sci U S A 102, 3184–3191.

Gray, J.M., Karow, D.S., Lu, H., Chang, A.J., Chang, J.S., Ellis, R.E., Marletta, M.A., and Bargmann, (2004). Oxygen sensation and social feeding mediated by a C. elegans guanylate cyclase homologue. Nature 430, 317–322.

Hawk, J.D., Calvo, A.C., Liu, P., Almoril-Porras, A., Aljobeh, A., Torruella-Suarez, M.L., Ren, I., Cook, N., Greenwood, J., Luo, L., et al. (2018). Integration of Plasticity Mechanisms within a Single Sensory Neuron of C. elegans Actuates a Memory. Neuron 97, 356-367.e354.

Hedgecock, E.M., and Russell, R.L. (1975). Normal and mutant thermotaxis in the nematode Caenorhabditis elegans. Proc Natl Acad Sci U S A 72, 4061–4065.

Hill, A.J., Mansfield, R., Lopez, J.M., Raizen, D.M., and Van Buskirk, C. (2014). Cellular stress induces a protective sleep-like state in C. elegans. Curr Biol 24, 2399–2405.

Hills, T., Brockie, P.J., and Maricq, A.V. (2004). Dopamine and glutamate control area-restricted search behavior in Caenorhabditis elegans. J Neurosci 24, 1217–1225.

Hostettler, L., Grundy, L., Kaser-Pebernard, S., Wicky, C., Schafer, W.R., and Glauser, D.A. (2017). The Bright Fluorescent Protein mNeonGreen Facilitates Protein Expression Analysis In Vivo. G3 (Bethesda) 7, 607–615.

Husson, S.J., Clynen, E., Baggerman, G., Janssen, T., and Schoofs, L. (2006). Defective processing of neuropeptide precursors in Caenorhabditis elegans lacking proprotein convertase 2 (KPC-2/EGL-3): mutant analysis by mass spectrometry. J Neurochem 98, 1999–2012.

Iannacone, M.J., Beets, I., Lopes, L.E., Churgin, M.A., Fang-Yen, C., Nelson, M.D., Schoofs, L., and Raizen, D.M. (2017). The RFamide receptor DMSR-1 regulates stress-induced sleep in C. elegans. Elife 6.

Iliff, A.J., Wang, C., Ronan, E.A., Hake, A.E., Guo, Y., Li, X., Zhang, X., Zheng, M., Liu, J., Grosh, K., et al. (2021). The nematode C. elegans senses airborne sound. Neuron 109, 3633–3646 e3637.

Ippolito, D., Thapliyal, S., and Glauser, D.A. (2021). Ca(2+)/CaM binding to CaMKI promotes IMA-3 importin binding and nuclear translocation in sensory neurons to control behavioral adaptation. Elife 10.

Javer, A., Currie, M., Lee, C.W., Hokanson, J., Li, K., Martineau, C.N., Yemini, E., Grundy, L.J., Li, C., Ch’ng, Q., et al. (2018a). An open-source platform for analyzing and sharing worm-behavior data. Nat Methods 15, 645–646.

Javer, A., Ripoll-Sanchez, L., and Brown, A.E.X. (2018b). Powerful and interpretable behavioural features for quantitative phenotyping of Caenorhabditis elegans. Philos Trans R Soc Lond B Biol Sci 373.

Ji, N., Madan, G.K., Fabre, G.I., Dayan, A., Baker, C.M., Kramer, T.S., Nwabudike, I., and Flavell, S.W. (2021). A neural circuit for flexible control of persistent behavioral states. Elife 10.

Jo, Y.S., Namboodiri, V.M.K., Stuber, G.D., and Zweifel, L.S. (2020). Persistent activation of central amygdala CRF neurons helps drive the immediate fear extinction deficit. Nat Commun 11, 422.

Jung, Y., Kennedy, A., Chiu, H., Mohammad, F., Claridge-Chang, A., and Anderson, D.J. (2020). Neurons that Function within an Integrator to Promote a Persistent Behavioral State in Drosophila. Neuron 105, 322–333 e325.

Kato, S., Kaplan, H.S., Schrodel, T., Skora, S., Lindsay, T.H., Yemini, E., Lockery, S., and Zimmer, M. (2015). Global brain dynamics embed the motor command sequence of Caenorhabditis elegans. Cell 163, 656–669.

Lea, S.E.G., Chow, P.K.Y., Leaver, L.A., and McLaren, I.P.L. (2020). Behavioral flexibility: A review, a model, and some exploratory tests. Learn Behav 48, 173–187.

Lee, R.Y., Sawin, E.R., Chalfie, M., Horvitz, H.R., and Avery, L. (1999). EAT-4, a homolog of a mammalian sodium-dependent inorganic phosphate cotransporter, is necessary for glutamatergic neurotransmission in caenorhabditis elegans. J Neurosci 19, 159–167.

Liu, P., Chen, B., and Wang, Z.W. (2020). GABAergic motor neurons bias locomotor decision-making in C. elegans. Nat Commun 11, 5076.

Lopez-Cruz, A., Sordillo, A., Pokala, N., Liu, Q., McGrath, P.T., and Bargmann, C.I. (2019). Parallel Multimodal Circuits Control an Innate Foraging Behavior. Neuron 102, 407–419 e408.

Magalhaes, A.C., Holmes, K.D., Dale, L.B., Comps-Agrar, L., Lee, D., Yadav, P.N., Drysdale, L., Poulter, M.O., Roth, B.L., Pin, J.P., et al. (2010). CRF receptor 1 regulates anxiety behavior via sensitization of 5-HT2 receptor signaling. Nat Neurosci 13, 622–629.

Marques, F., Saro, G., Lia, A.S., Poole, R.J., Falquet, L., and Glauser, D.A. (2019). Identification of avoidance genes through neural pathway-specific forward optogenetics. PLoS Genet 15, e1008509.

Marquina-Solis, J., Vandewyer, E., Hawk, J., Colón-Ramos, D.A., Beets, I., and Bargmann, C.I. (2022). Peptidergic signaling controls the dynamics of sickness behavior in <em>Caenorhabditis elegans</em>. bioRxiv, 2022.2004.2016.488560.

Martineau, C.N., Brown, A.E.X., and Laurent, P. (2020). Multidimensional phenotyping predicts lifespan and quantifies health in Caenorhabditis elegans. PLoS Comput Biol 16, e1008002.

Matsumoto, M., Straub, R.E., Marenco, S., Nicodemus, K.K., Matsumoto, S., Fujikawa, A., Miyoshi, S., Shobo, M., Takahashi, S., Yarimizu, J., et al. (2008). The evolutionarily conserved G protein-coupled receptor SREB2/GPR85 influences brain size, behavior, and vulnerability to schizophrenia. Proc Natl Acad Sci U S A 105, 6133–6138.

Metsalu, T., and Vilo, J. (2015). ClustVis: a web tool for visualizing clustering of multivariate data using Principal Component Analysis and heatmap. Nucleic Acids Res 43, W566–570.

Miyabayashi, T., Palfreyman, M.T., Sluder, A.E., Slack, F., and Sengupta, P. (1999). Expression and function of members of a divergent nuclear receptor family in Caenorhabditis elegans. Dev Biol 215, 314–331.

Nichols, A.L.A., Eichler, T., Latham, R., and Zimmer, M. (2017). A global brain state underlies C. elegans sleep behavior. Science 356.

Ohnishi, N., Kuhara, A., Nakamura, F., Okochi, Y., and Mori, I. (2011). Bidirectional regulation of thermotaxis by glutamate transmissions in Caenorhabditis elegans. EMBO J 30, 1376–1388.

Oranth, A., Schultheis, C., Tolstenkov, O., Erbguth, K., Nagpal, J., Hain, D., Brauner, M., Wabnig, S., Steuer Costa, W., McWhirter, R.D., et al. (2018). Food Sensation Modulates Locomotion by Dopamine and Neuropeptide Signaling in a Distributed Neuronal Network. Neuron 100, 1414–1428 e1410.

Raizen, D.M., Zimmerman, J.E., Maycock, M.H., Ta, U.D., You, Y.J., Sundaram, M.V., and Pack, A.I. (2008). Lethargus is a Caenorhabditis elegans sleep-like state. Nature 451, 569–U566.

Ringstad, N., and Horvitz, H.R. (2008). FMRFamide neuropeptides and acetylcholine synergistically inhibit egg-laying by C. elegans. Nat Neurosci 11, 1168–1176.

Saro, G., Lia, A.S., Thapliyal, S., Marques, F., Busch, K.E., and Glauser, D.A. (2020). Specific Ion Channels Control Sensory Gain, Sensitivity, and Kinetics in a Tonic Thermonociceptor. Cell Rep 30, 397–408 e394.

Schild, L.C., and Glauser, D.A. (2013). Dynamic switching between escape and avoidance regimes reduces Caenorhabditis elegans exposure to noxious heat. Nat Commun 4, 2198.

Schild, L.C., and Glauser, D.A. (2015). Dual Color Neural Activation and Behavior Control with Chrimson and CoChR in Caenorhabditis elegans. Genetics 200, 1029–1034.

Schild, L.C., Zbinden, L., Bell, H.W., Yu, Y.V., Sengupta, P., Goodman, M.B., and Glauser, D.A. (2014). The balance between cytoplasmic and nuclear CaM kinase-1 signaling controls the operating range of noxious heat avoidance. Neuron 84, 983–996.

Schmitt, C., Schultheis, C., Pokala, N., Husson, S.J., Liewald, J.F., Bargmann, C.I., and Gottschalk, A. (2012). Specific expression of channelrhodopsin-2 in single neurons of Caenorhabditis elegans. PLoS One 7, e43164.

Shaw, P.J., Tononi, G., Greenspan, R.J., and Robinson, D.F. (2002). Stress response genes protect against lethal effects of sleep deprivation in Drosophila. Nature 417, 287–291.

Shtonda, B.B., and Avery, L. (2006). Dietary choice behavior in Caenorhabditis elegans. J Exp Biol 209, 89–102.

Skora, S., Mende, F., and Zimmer, M. (2018). Energy Scarcity Promotes a Brain-wide Sleep State Modulated by Insulin Signaling in C. elegans. Cell Rep 22, 953–966.

Sorrells, T.R., Pandey, A., Rosas-Villegas, A., and Vosshall, L.B. (2022). A persistent behavioral state enables sustained predation of humans by mosquitoes. Elife 11.

Stern, S., Kirst, C., and Bargmann, C.I. (2017). Neuromodulatory Control of Long-Term Behavioral Patterns and Individuality across Development. Cell 171, 1649–1662 e1610.

Swierczek, N.A., Giles, A.C., Rankin, C.H., and Kerr, R.A. (2011). High-throughput behavioral analysis in C. elegans. Nat Methods 8, 592–598.

Taylor, S.R., Santpere, G., Weinreb, A., Barrett, A., Reilly, M.B., Xu, C., Varol, E., Oikonomou, P., Glenwinkel, L., McWhirter, R., et al. (2021). Molecular topography of an entire nervous system. Cell 184, 4329-4347.e4323.

Uddin, L.Q. (2021). Cognitive and behavioural flexibility: neural mechanisms and clinical considerations. Nat Rev Neurosci 22, 167–179.

Venkatachalam, V., Ji, N., Wang, X., Clark, C., Mitchell, J.K., Klein, M., Tabone, C.J., Florman, J., Ji, H., Greenwood, J., et al. (2016). Pan-neuronal imaging in roaming Caenorhabditis elegans. Proc Natl Acad Sci U S A 113, E1082–1088.

Waggoner, L.E., Zhou, G.T., Schafer, R.W., and Schafer, W.R. (1998). Control of alternative behavioral states by serotonin in Caenorhabditis elegans. Neuron 21, 203–214.

Wakabayashi, T., Kitagawa, I., and Shingai, R. (2004). Neurons regulating the duration of forward locomotion in Caenorhabditis elegans. Neurosci Res 50, 103–111.

Xiao, R., and Xu, X.Z.S. (2021). Temperature Sensation: From Molecular Thermosensors to Neural Circuits and Coding Principles. Annu Rev Physiol 83, 205–230.

Yemini, E., Jucikas, T., Grundy, L.J., Brown, A.E., and Schafer, W.R. (2013). A database of Caenorhabditis elegans behavioral phenotypes. Nat Methods 10, 877–879.

