## Supplementary Figures for "Multisite gating in tonic sensory circuits integrates multimodal context to control persistent behavioral states"

### SUPPLEMENTAL INFORMATION

**Figure S1.1. Recording device, experimental pipeline overview and list of extracted behavioral parameters for motion and posture.**

Schematic of the recording device used for behavioral recording in a tightly controlled thermal environment (**A**). The whole system is inside an incubator and isolated from the intrinsic vibration generated by the device via an anti-vibration platform (note that both the camera and the recorded plate are hence isolated). Overview of the experimental pipeline (**B**) and of the extracted Tierpsy tracker parameters (**C**).

**Figure S1.2. Behavioral codes during food and temperature-dependent behavioral transitions**

Heat-map of behavioral parameters of *z*-scores across the indicated conditions and hierarchical clustering based on Euclidian distance (tree on the left). Clusters of parameters similarly affected by starvation, growth temperature and/or recent temperature shift are annotated (colored labels on the right). A time series over 6 h is presented for each condition based on the same data set as the one use for PCA analyses reported in Figure 1. Each data point represents the average value for 1-min recordings on at least three independent worm populations (>40 worm each).

**Figure S1.3. Temperature and food effects on behavioral states based on motion behavioral parameters**

Same analyses and displays as in Figure 1, but using only the motion parameters as defined in Fig S1.1C. Panel M is a summary of the steady states after 6 h, and is also presented in Fig. 1M.

**Figure S1.4. Temperature and food effects on behavioral states based on postural behavioral parameters**

Same analyses and displays as in Figure 1, but using only the postural parameters as defined in Fig S1.1C. Panel M is a summary of the steady states after 6 h, and is also presented in Fig. 1N.

**Figure S2. Dispersal trajectory simulations comparing dwelling, global and glocal search states**

Results of Monte-Carlo simulations considering the average frequency of turns and speed measured in worm populations in isothermal environments. Thirty-five 1-min trajectories for each condition (**A**). Average (±s.e.m) and individual data points for animal displacement (**B**, corresponding to how far animals moved from their starting point) and covered distance (**C**, corresponding to the path length of each track). Both simulated and real data are presented side-by-side. Real data are the same as in Fig. 2.

**Figure S4. Postural alteration during transition to scanning depends on AFD neuron, FLP-6 neuropeptide and multiple GPCRs**

Time course of worm posture alteration (decrease in tail bending) after warming from 15 to 25°C (**A**). Tail bend at $t$=0 and $t$=6 h following warming (thermal shift from 15 to 25°C) in the indicated transgenic or mutant strains (**B**, **C**, **D**). Bars as mean ± s.e.m. of $n \geq 5$ assays each with ≥50 worms. *Pgcy-8::TeTx*, transgene blocking synaptic transmission in AFD (**B**); *Pgcy-8::flp-6,* transgene for AFD-specific *flp-6* rescue (**C**); *Pegl-6::egl-6*, transgene for *egl-6* over-expression and *flp-6* mutation by-pass analyses in *egl-6* and *flp-6* mutant background, respectively (**D**). **, $p<.01$ versus 15°C Fed condition, # $p<.05$ and ##, $p<.01$ versus the indicated control by Bonferroni posthoc tests.

**Figure S5. FLP thermosensory neurons are essential for omega turn increase during glocal search**

Time course of omega turn frequency increase after starvation at 15 or 25°C (15 Starved, 25 Starved) and control left on food at respective temperatures (15 Fed, 25 Fed) (**A**). Mean ± s.e.m. of *n*=6 assays each with ≥50 worms. Omega turn frequency measured after 6h of starvation at 25°C in wild type (N2), in transgenic lines with genetic ablation of the indicated neurons, or in animals carrying a *Pmec-3::TeTx* transgene blocking neurotransmission in FLP (**B**). Like for speed elevation (Fig. 5), FLP plays a major role in the up-regulation of omega turns.

**Figure S6. FLP thermosensory neurons are essential for omega turn increase during glocal search**

Impact of a *cat*-2 mutation blocking dopamine biosynthesis on the omega turn frequency in fed animals at 15 or 25°C (**A**). Impact of *eat-4* mutation affecting glutamatergic signaling in starved animals at 15 (blue) or 25°C (orange) (**B**). Mean ± s.e.m. of $n \geq 7$ assays, each scoring ≥30 worms. **, $p$<.01 versus N2; # $p$<.05 and ##, $p$<.01 versus the indicated condition by Bonferroni posthoc tests. ns, not significant.

**Figure S7. Omega turn up-regulation during glocal search involves FLP-5/DMSR-1 signaling from FLP**

Genetic dissection of the molecular signaling controlling omega turn frequency increase during glocal search. Impact of neuropeptide-affecting mutations on the omega turn frequency of starved animals held at 25°C (**A**). Impact of *flp-5* mutation, over-expression with a *Pflp-5::flp-5* transgene, and rescue/over-expression with *Pmec-3::flp-5* transgene expressed in FLP (**B**). Impact of mutations affecting FLP-5 and its receptors (**C**). Impact of gain-of-function (gf) and loss-of-function (lf) mutations in *egl-6*, as well as FLP-5 receptor over-expression in *egl-6*-expressing cells, revealing that EGL-6, DMSR-1a, DMSR-1b and DMSR-7 have a similar inhibitory effect on omega turn frequency in starved animals at 25°C (**D**). DMSR-1 overexpression in *dmsr-1*-expressing cells or AVA-specific overexpression, respectively, on the speed of starved animals at 25°C (**E**).

**Figure S8. Model**

**A** Temperature-controlled recording device

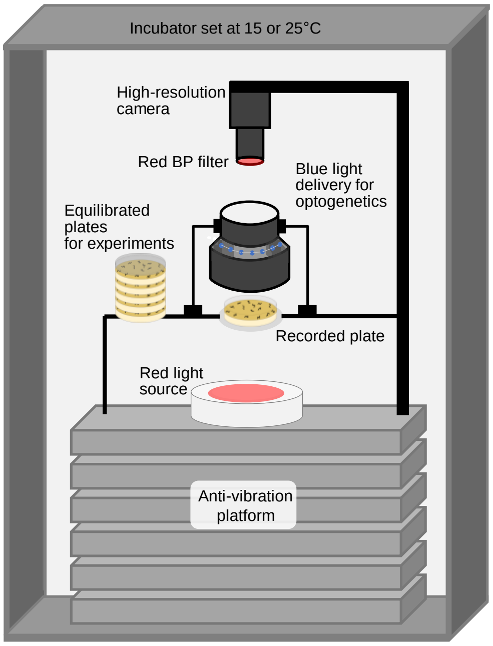


**B** Experimental pipeline overview

#### Worm preparation

- Worm synchronization by bleaching
- Growth on regular NGM plate seeded with *E. coli* at 15°C or 25°C with adjusted growth time until young adult stage.
- For starvation, worm washing and return to a new food-free plate.

#### Recording

- Transfer of plates in the recording device pre-equilibrated at the target recording temperature (15 or 25°C)
- High-resolution movie recordings (3 min) (optional: optogenetic stimulation)

#### Data analysis:

- Batch processing of movie using Tierpsy tracker
- Extraction of behavioral features
- PCA and hierarchical clustering using ClustVis
- Specific analyses on selected parameters

**C** Extracted Tierpsy parameters

**Motion**

**crawling amplitude**
- head_crawling_amplitude_abs
- midbody_crawling_amplitude_abs
- tail_crawling_amplitude_abs

**crawling frequency**
- head_crawling_frequency_abs
- midbody_crawling_frequency_abs
- tail_crawling_frequency_abs

**foraging**
- foraging_speed_abs
- foraging_amplitude_abs

**path**
- path_range
- path_curvature_abs

**forward locomotion**
- forward_time
- inter_forward_time
- forward_frequency
- forward_time_ratio

**backward locomotion**
- backward_time
- inter_backward_time
- backward_frequency
- backward_time_ratio

**speed of locomotion**
- midbody_speed_abs

**dwelling**
- midbody_dwelling

**omega turns**
- omega_turns_time
- inter_omega_turns_time
- omega_turns_frequency
- omega_turns_time_ratio

**pausing**
- paused_time
- inter_paused_time
- paused_frequency
- paused_time_ratio

**forward speed / length**
- forward_speed_length_normalized

**angular speed**
- midbody_motion_direction_abs

**Posture**

**body bend**
- bend count
- head_bend_mean_abs
- neck_bend_mean_abs
- midbody_bend_mean_abs
- hips_bend_mean_abs
- tail_bend_mean_abs

**body bend variability**
- head_bend_sd_abs
- neck_bend_sd_abs
- midbody_bend_sd_abs
- hips_bend_sd_abs
- tail_bend_sd_abs

**eigen projections**
- eigen_projection_1_abs
- eigen_projection_2_abs
- eigen_projection_3_abs
- eigen_projection_4_abs
- eigen_projection_5_abs
- eigen_projection_6_abs

**Figure S1.1**

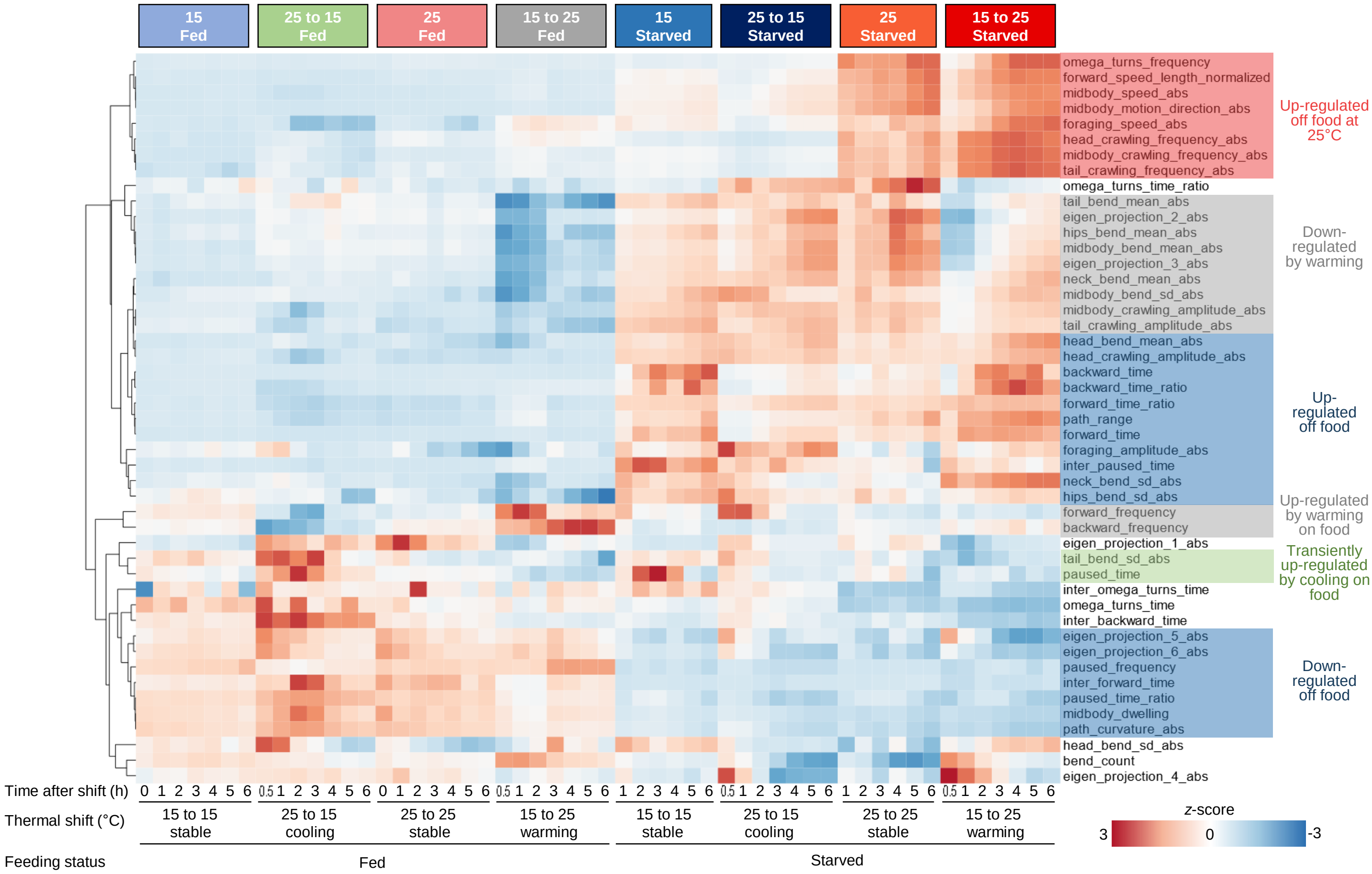
15 Fed
25 to 15 Fed
25 Fed
15 to 25 Fed
15 Starved
25 to 15 Starved
25 Starved
15 to 25 Starved
omega_turns_frequency
forward_speed_length_normalized
midbody_speed_abs
midbody_motion_direction_abs
foraging_speed_abs
head_crawling_frequency_abs
midbody_crawling_frequency_abs
tail_crawling_frequency_abs
omega_turns_time_ratio
tail_bend_mean_abs
eigen_projection_2_abs
hips_bend_mean_abs
midbody_bend_mean_abs
eigen_projection_3_abs
neck_bend_mean_abs
midbody_bend_sd_abs
midbody_crawling_amplitude_abs
tail_crawling_amplitude_abs
head_bend_mean_abs
head_crawling_amplitude_abs
backward_time
backward_time_ratio
forward_time_ratio
path_range
forward_time
foraging_amplitude_abs
inter_paused_time
neck_bend_sd_abs
hips_bend_sd_abs
forward_frequency
backward_frequency
eigen_projection_1_abs
tail_bend_sd_abs
paused_time
inter_omega_turns_time
omega_turns_time
inter_backward_time
eigen_projection_5_abs
eigen_projection_6_abs
paused_frequency
inter_forward_time
paused_time_ratio
midbody_dwelling
path_curvature_abs
head_bend_sd_abs
bend_count
eigen_projection_4_abs
Up-regulated off food at 25°C
Down-regulated by warming
Up-regulated off food
Up-regulated by warming on food
Transiently up-regulated by cooling on food
Down-regulated off food
Time after shift (h)
0 1 2 3 4 5 6
0.5 1 2 3 4 5 6
0 1 2 3 4 5 6
0.5 1 2 3 4 5 6
1 2 3 4 5 6
0.5 1 2 3 4 5 6
1 2 3 4 5 6
0.5 1 2 3 4 5 6
Thermal shift (°C)
15 to 15 stable
25 to 15 cooling
25 to 25 stable
15 to 25 warming
15 to 15 stable
25 to 15 cooling
25 to 25 stable
15 to 25 warming
Feeding status
Fed
Starved
z-score
3
0
-3

**Figure S1.2**

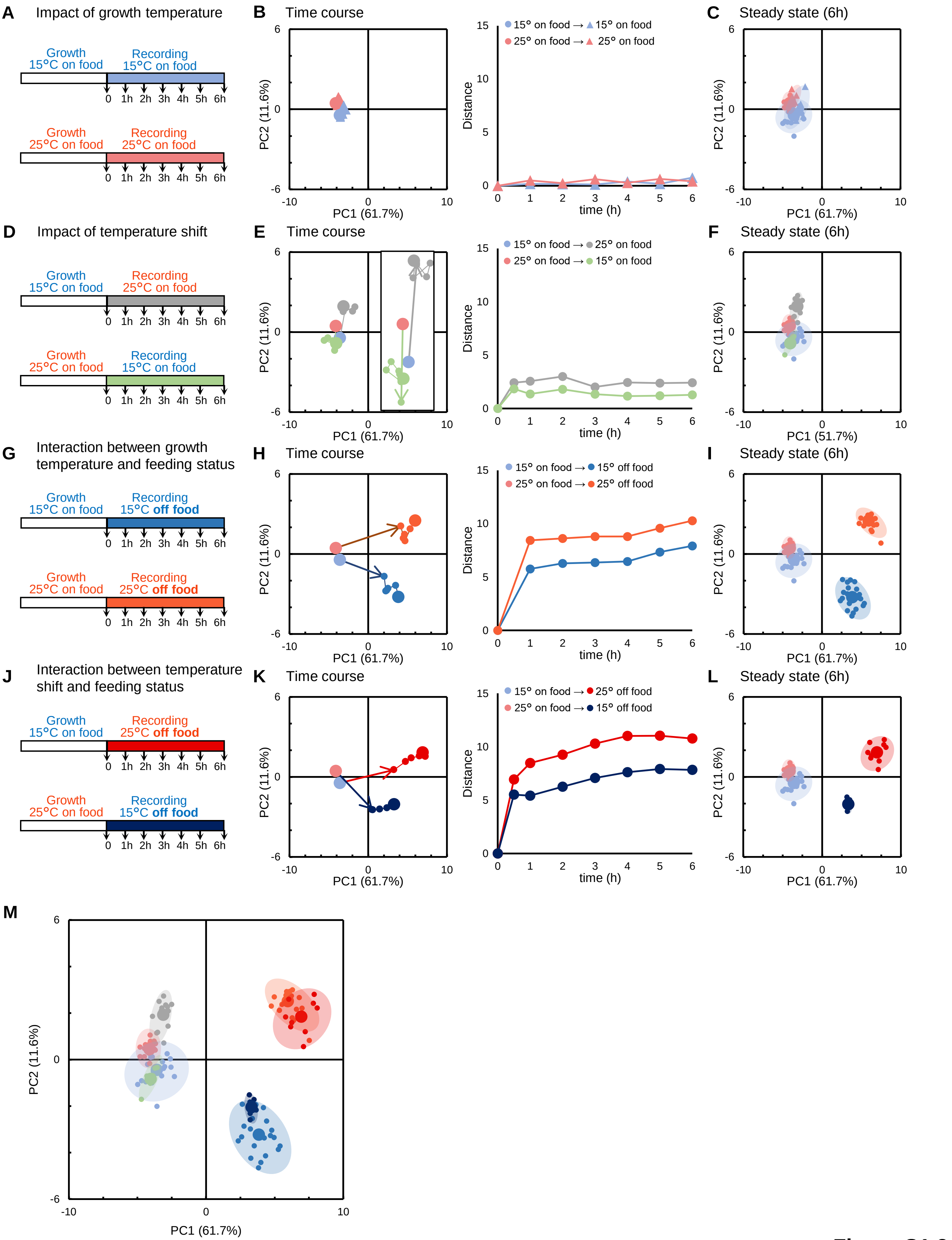
A Impact of growth temperature
Growth 15°C on food
Recording 15°C on food
Growth 25°C on food
Recording 25°C on food
0 1h 2h 3h 4h 5h 6h
B Time course
PC1 (61.7%)
PC2 (11.6%)
15° on food → 15° on food
25° on food → 25° on food
Distance
time (h)
C Steady state (6h)
D Impact of temperature shift
Growth 15°C on food
Recording 25°C on food
Growth 25°C on food
Recording 15°C on food
E Time course
15° on food → 25° on food
25° on food → 15° on food
F Steady state (6h)
PC1 (51.7%)
G Interaction between growth temperature and feeding status
Growth 15°C on food
Recording 15°C off food
Growth 25°C on food
Recording 25°C off food
H Time course
15° on food → 15° off food
25° on food → 25° off food
I Steady state (6h)
J Interaction between temperature shift and feeding status
Growth 15°C on food
Recording 25°C off food
Growth 25°C on food
Recording 15°C off food
K Time course
15° on food → 25° off food
25° on food → 15° off food
L Steady state (6h)
M

**Figure S1.3**

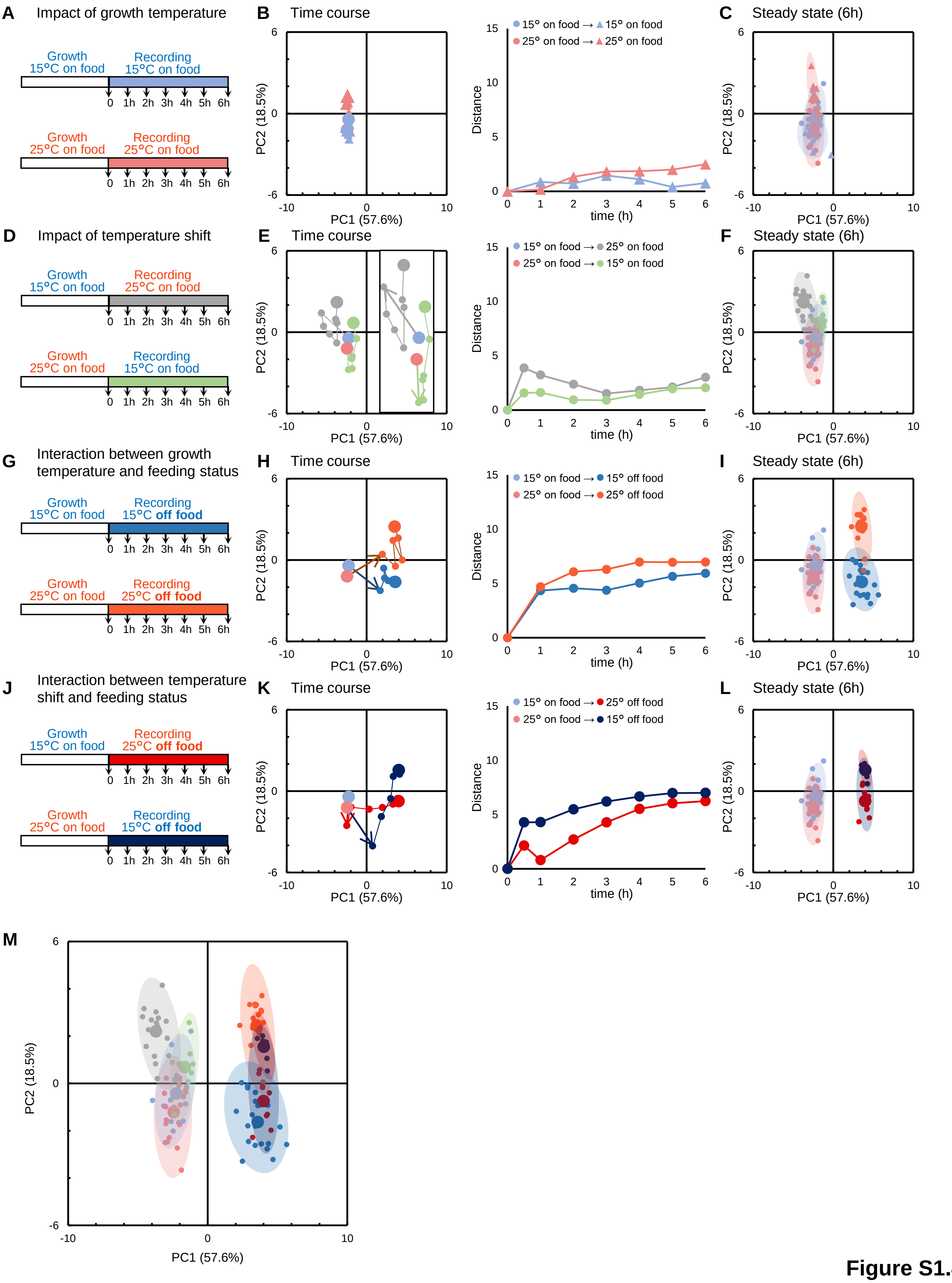
A Impact of growth temperature
Growth 15°C on food
Recording 15°C on food
Growth 25°C on food
Recording 25°C on food
0 1h 2h 3h 4h 5h 6h
B Time course
15° on food → 15° on food
25° on food → 25° on food
C Steady state (6h)
D Impact of temperature shift
Growth 15°C on food
Recording 25°C on food
Growth 25°C on food
Recording 15°C on food
E Time course
15° on food → 25° on food
25° on food → 15° on food
F Steady state (6h)
G Interaction between growth temperature and feeding status
Growth 15°C on food
Recording 15°C off food
Growth 25°C on food
Recording 25°C off food
H Time course
15° on food → 15° off food
25° on food → 25° off food
I Steady state (6h)
J Interaction between temperature shift and feeding status
Growth 15°C on food
Recording 25°C off food
Growth 25°C on food
Recording 15°C off food
K Time course
15° on food → 25° off food
25° on food → 15° off food
L Steady state (6h)
M
PC1 (57.6%)
PC2 (18.5%)
Distance
time (h)

Figure S1.4

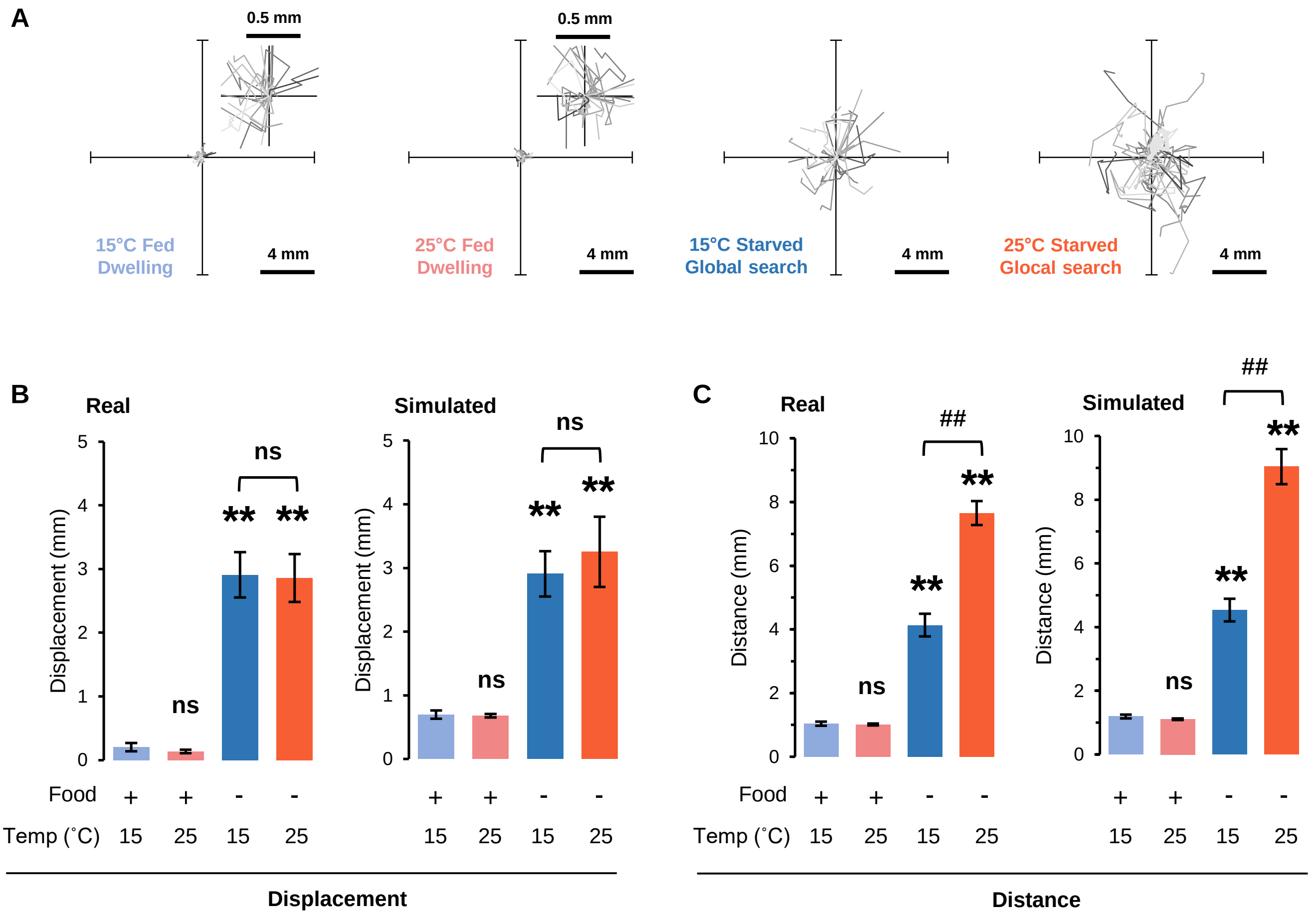

A
0.5 mm
0.5 mm
15°C Fed
Dwelling
4 mm
25°C Fed
Dwelling
4 mm
15°C Starved
Global search
4 mm
25°C Starved
Glocal search
4 mm
B
Real
Simulated
Displacement (mm)
ns
**
ns
C
Real
Simulated
Distance (mm)
##
**
ns
Food + + - -
Temp (˚C) 15 25 15 25
Displacement
Distance


Figure S2

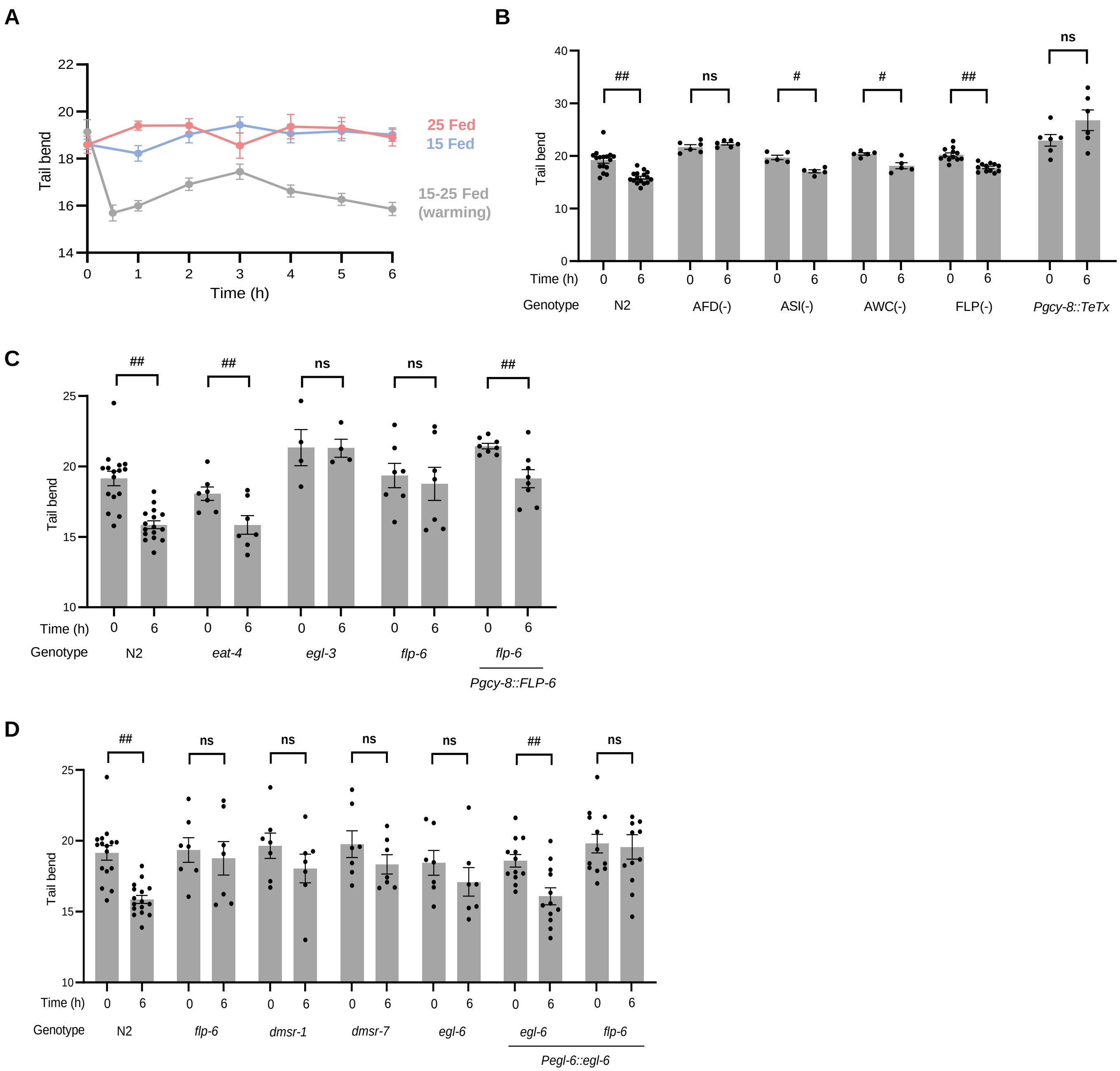
A
Tail bend
Time (h)
25 Fed
15 Fed
15-25 Fed (warming)
B
Tail bend
Time (h)
Genotype
N2
AFD(-)
ASI(-)
AWC(-)
FLP(-)
Pgcy-8::TeTx
##
ns
#
C
Tail bend
Time (h)
Genotype
N2
eat-4
egl-3
flp-6
flp-6
Pgcy-8::FLP-6
D
Tail bend
Time (h)
Genotype
N2
flp-6
dmsr-1
dmsr-7
egl-6
egl-6
flp-6
Pegl-6::egl-6

**Figure S4**

A

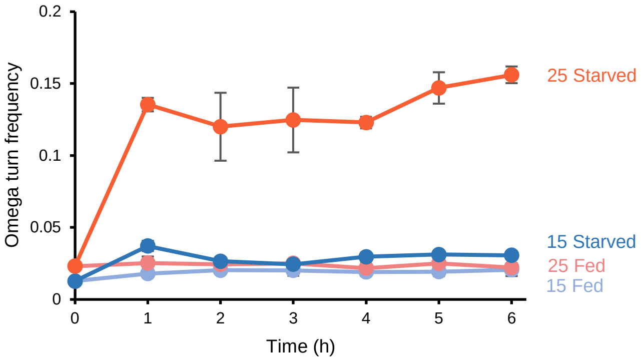

Omega turn frequency
0.2
0.15
0.1
0.05
0
0
1
2
3
4
5
6
Time (h)
25 Starved
15 Starved
25 Fed
15 Fed


B

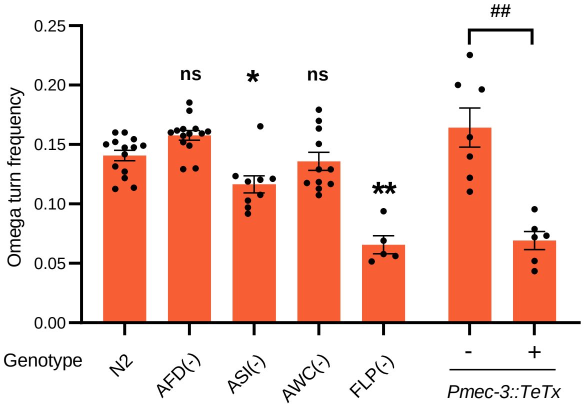

##
ns
*
ns
**
Omega turn frequency
0.25
0.20
0.15
0.10
0.05
0.00
Genotype
N2
AFD(-)
ASI(-)
AWC(-)
FLP(-)
-
+
Pmec-3::TeTx


**Figure S5**

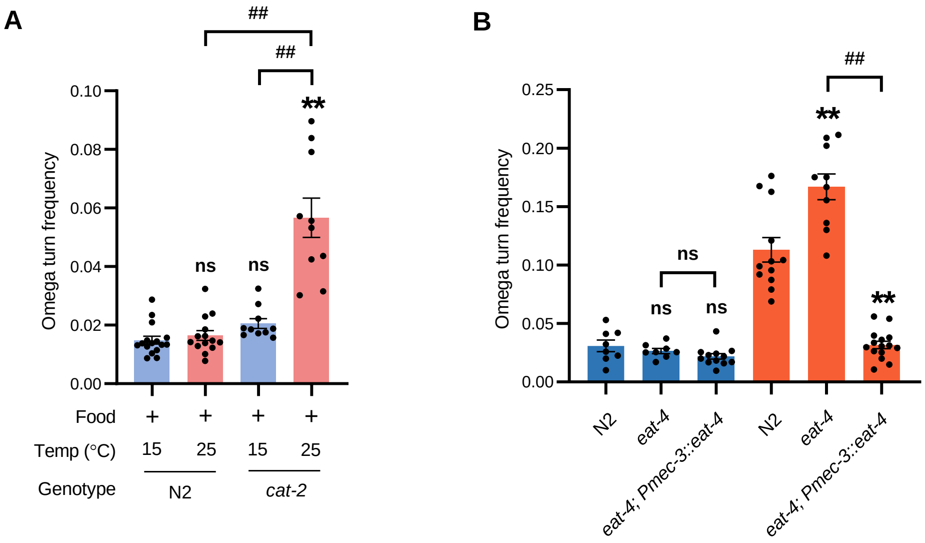
A
##
##
**
ns
ns
Omega turn frequency
0.10
0.08
0.06
0.04
0.02
0.00
Food
+
+
+
+
Temp (°C)
15
25
15
25
Genotype
N2
cat-2
B
##
**
ns
ns
ns
**
Omega turn frequency
0.25
0.20
0.15
0.10
0.05
0.00
N2
eat-4
eat-4; Pmec-3::eat-4
N2
eat-4
eat-4; Pmec-3::eat-4

**Figure S6**

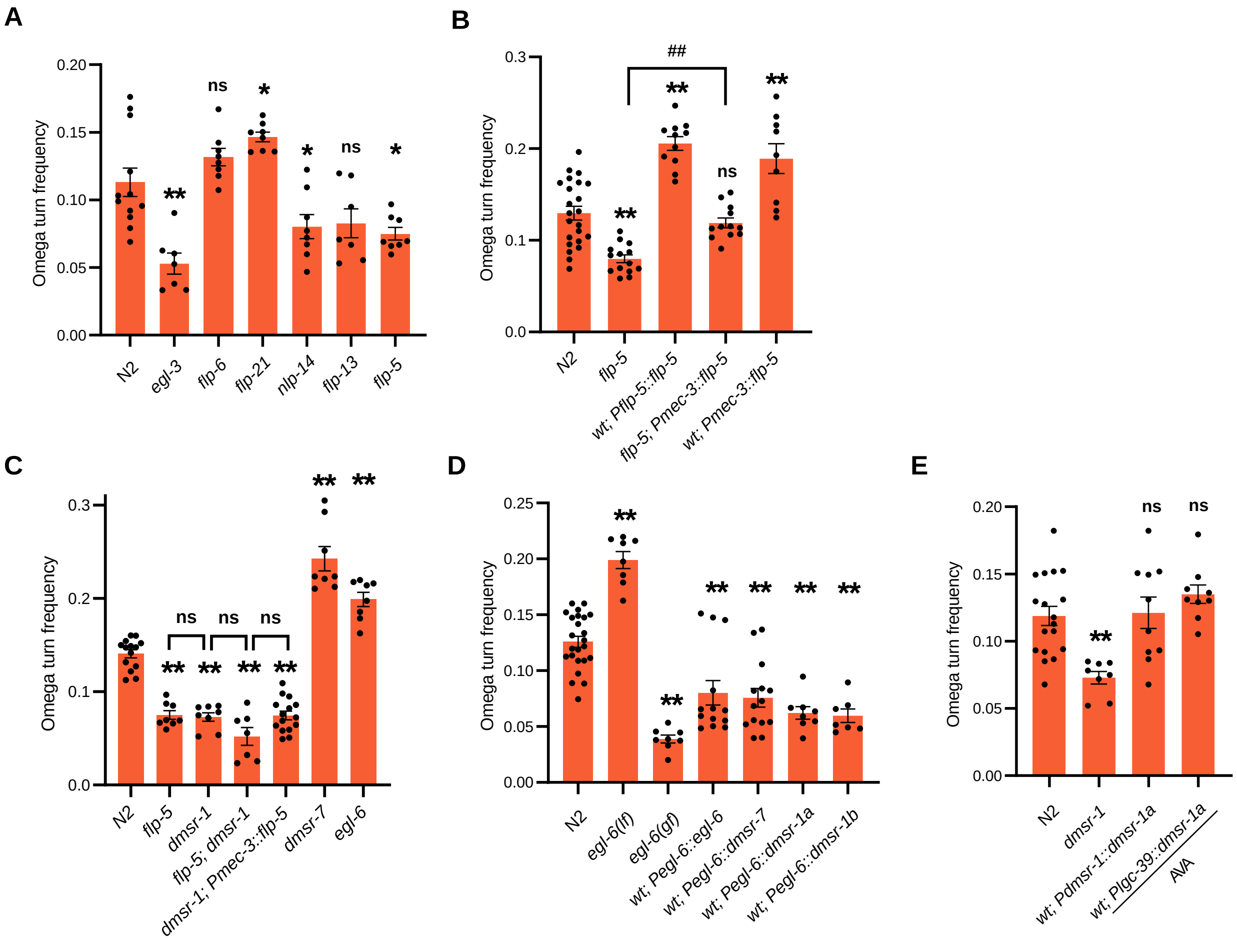

A
Omega turn frequency
0.00
0.05
0.10
0.15
0.20
**
ns
*
*
ns
*
N2
egl-3
flp-6
flp-21
nlp-14
flp-13
flp-5
B
Omega turn frequency
0.0
0.1
0.2
0.3
##
**
**
ns
**
N2
flp-5
wt; Pflp-5::flp-5
flp-5; Pmec-3::flp-5
wt; Pmec-3::flp-5
C
Omega turn frequency
0.0
0.1
0.2
0.3
ns
ns
ns
**
**
**
**
**
**
N2
flp-5
dmsr-1
flp-5; dmsr-1
dmsr-1; Pmec-3::flp-5
dmsr-7
egl-6
D
Omega turn frequency
0.00
0.05
0.10
0.15
0.20
0.25
**
**
**
**
**
**
N2
egl-6(lf)
egl-6(gf)
wt; Pegl-6::egl-6
wt; Pegl-6::dmsr-7
wt; Pegl-6::dmsr-1a
wt; Pegl-6::dmsr-1b
E
Omega turn frequency
0.00
0.05
0.10
0.15
0.20
**
ns
ns
N2
dmsr-1
wt; Pdmsr-1::dmsr-1a
wt; Pglc-39::dmsr-1a
AVA


Figure S7

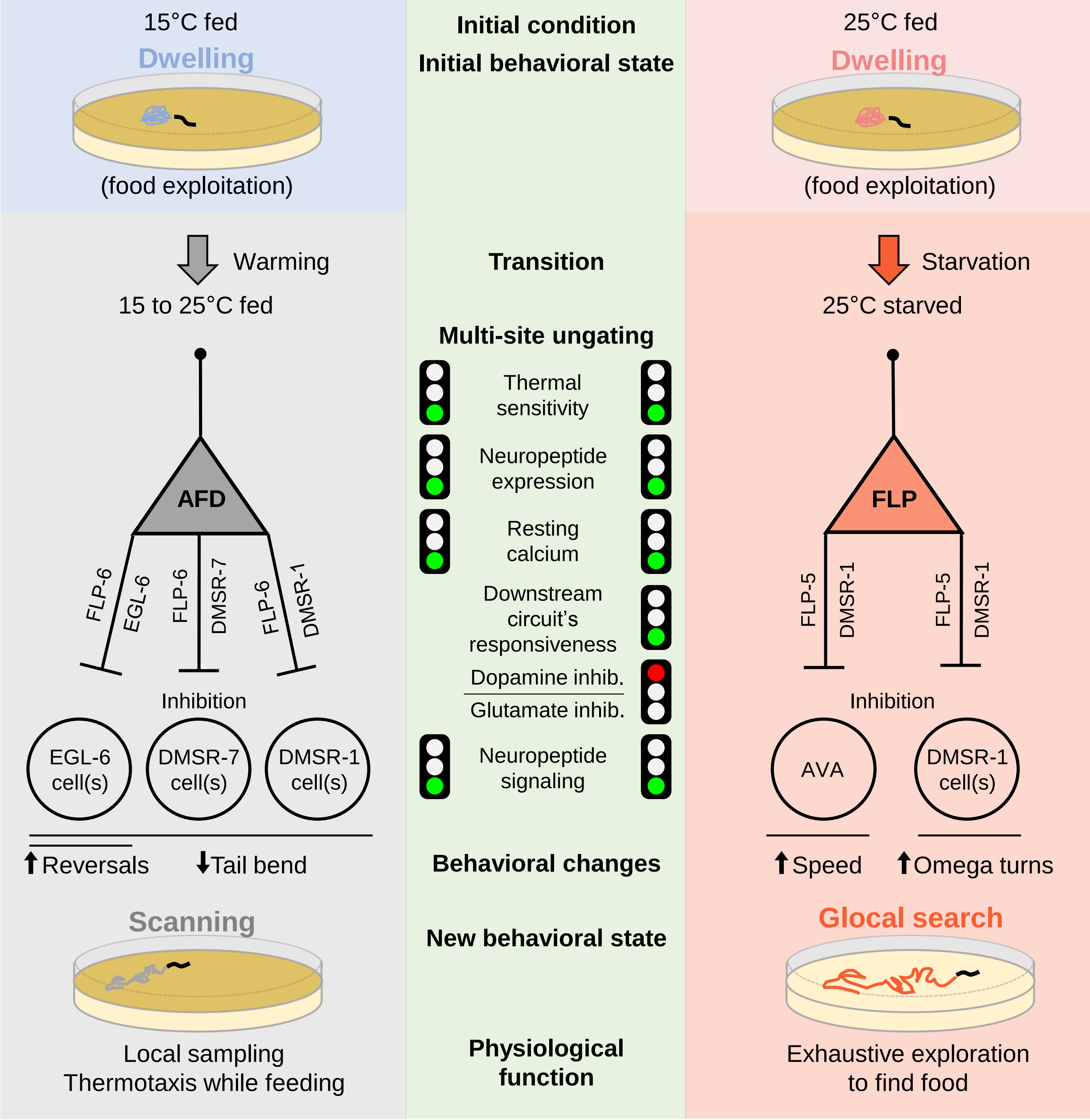

15°C fed
Dwelling
(food exploitation)
Initial condition
Initial behavioral state
25°C fed
Dwelling
(food exploitation)
Warming
15 to 25°C fed
Transition
Starvation
25°C starved
Multi-site ungating
Thermal sensitivity
Neuropeptide expression
Resting calcium
Downstream circuit's responsiveness
Dopamine inhib.
Glutamate inhib.
Neuropeptide signaling
AFD
FLP-6
EGL-6
FLP-6
DMSR-7
FLP-6
DMSR-1
Inhibition
EGL-6 cell(s)
DMSR-7 cell(s)
DMSR-1 cell(s)
FLP
FLP-5
DMSR-1
FLP-5
DMSR-1
Inhibition
AVA
DMSR-1 cell(s)
↑Reversals
↓Tail bend
Behavioral changes
↑Speed
↑Omega turns
Scanning
New behavioral state
Glocal search
Local sampling
Thermotaxis while feeding
Physiological function
Exhaustive exploration to find food


**Figure S8**
